# Multiplex, translaminar imaging in the spinal cord of behaving mice

**DOI:** 10.1101/2021.12.23.474039

**Authors:** Pavel Shekhtmeyster, Erin M. Carey, Daniela Duarte, Alexander Ngo, Grace Gao, Nicholas A. Nelson, Charles L. Clark, Axel Nimmerjahn

**Affiliations:** Waitt Advanced Biophotonics Center, The Salk Institute for Biological Studies, La Jolla, CA 92037, USA; Electrical and Computer Engineering Graduate Program, University of California, San Diego, La Jolla, CA 92037, USA; Biological Sciences Graduate Program, University of California, San Diego, La Jolla, CA 92037, USA

## Abstract

While the spinal cord is known to play critical roles in sensorimotor processing, including pain-related signaling, corresponding activity patterns in genetically defined cell types across spinal laminae have remained elusive. Calcium imaging has enabled cellular activity measurements in behaving rodents but is currently limited to superficial regions. Using chronically implanted microprisms, we imaged sensory and motor evoked activity in regions and at speeds inaccessible by other high-resolution imaging techniques. To enable translaminar imaging in freely behaving animals through implanted microprisms, we additionally developed wearable microscopes with custom-compound microlenses. This new integrated system addresses multiple challenges of previous wearable microscopes, including their limited working distance, resolution, contrast, and achromatic range. The combination of these innovations allowed us to uncover that dorsal horn astrocytes in behaving mice show somatosensory program-dependent and lamina-specific calcium excitation. Additionally, we show that tachykinin precursor 1 (Tac1)-expressing neurons exhibit upper laminae-restricted activity to acute mechanical pain but not locomotion.

## Introduction

The spinal cord plays crucial roles in somatosensation^1–3^. Genetic, pharmacologic, anatomical, and electrophysiological studies have revealed that specific somatosensory programs, such as pain or movement, are linked to spatially restricted modules comprised of heterogeneous neuronal cell types. These modules reside in defined laminae, located at different depths from the spinal surface^1–3^. However, genetic and pharmacologic approaches lack the spatial and temporal resolution to investigate cellular activity patterns that underlie somatosensory computations and typically rely on overt behavioral phenotypes to define cell types’ functional roles. Histological methods can identify highly active neurons across spinal laminae (e.g., cFos expression) but generate only snapshots at discrete time points. Electrophysiology approaches are valuable for probing neuronal activity, but recordings in behaving animals from genetically defined cells across spinal laminae remain to be demonstrated. Emerging evidence also suggests that spinal astrocytes respond to different kinds of neural activity with calcium excitation and modulate pain signaling and motor function^4–8^. Astrocyte activity remains difficult to interrogate with the methods mentioned above.

Fluorescence imaging has recently allowed real-time measurement of neuronal and astrocyte activity in the spinal cord of behaving mice^4^. However, scattering and absorption restrict optical access to superficial dorsal horn regions. This is because the spinal gray matter is surrounded by thick myelin layers that are highly reflective across a wide wavelength range. This anatomical arrangement limits one- and two-photon imaging to about 75-150 μm and 150-350 μm depths, respectively, depending on the fluorescence indicator employed^9^. With three-photon microscopy, depths of about 350-500 μm can be achieved, but at the expense of acquisition frame rate, the field of view (FOV), or multi-color imaging capabilities^10^. Low acquisition frame rates are particularly detrimental to spinal cord recordings, as they hamper computational correction of tissue movement-induced image artifacts. The large-amplitude and non-uniform tissue displacements typical during animal behavior are also the reason why electrophysiological recordings regularly used in other central nervous system regions have proven extremely challenging in the spinal cord^9^.

The implantation of micro-optics, such as gradient-index (GRIN) lenses, has allowed access to deep tissue regions in the brain. However, typical GRIN lens diameters (≥0.5 mm) are comparable to the spinal gray matter’s width in mice, preventing their insertion without undue tissue disruption. In contrast, glass reflective microprisms have enabled optical access to brain areas difficult if not impossible to reach otherwise, such as prefrontal and entorhinal cortex, and without tissue removal^11–13^. Whether and how a microprism-based approach could be applied to the spinal cord remains unclear.

Imaging through implanted microprisms requires long working distance (WD) objectives. Current wearable microscopes for imaging in freely behaving animals do not offer WDs >1 mm and often rely on GRIN objectives, preventing high-resolution imaging through these micro-optics^9, 14, 15^. The reliance on GRIN-based imaging optics also limits multi-color recording capabilities due to their narrow achromatic range (tens of nanometers)^16^.

These technical barriers in both surgery and imaging have prevented measurement of cellular activity in deep sensory and premotor areas and at high speed across spinal laminae of behaving animals, limiting our ability to uncover how the spatiotemporal activity patterns in genetically defined cell types contribute to sensorimotor processing. To address these long-standing challenges, we introduce several innovations, including 1) a wearable microscope with custom-compound-microlenses allowing high-speed (45 fps), high-resolution and high-contrast (∼1.5 μm), and multi-color imaging (500-620 nm spectral range) through tissue-implanted micro-optics (2.4-mm working distance), and 2) a surgical procedure that enables long-term activity measurements across spinal laminae using implanted microprisms. Using these methods, we show that tachykinin precursor 1 (Tac1)-expressing neurons respond rapidly to painful mechanical stimuli specifically in sensory dorsal horn laminae. In contrast, astrocytes showed slow calcium transients to the same stimuli in sensory and motor-evoked activity in premotor areas, suggesting region-dependent (i.e., lamina- and neuronal cell type-dependent) functional signaling within this glial cell population.

## Results

### Wearable microscopes with long working distance custom-compound-microlenses for high-resolution, high-contrast measurements in freely behaving mice

To meet the application needs mentioned above, as well as match or exceed the optical performance characteristics of comparable wearable devices, the design goals for our microscope’s optical system included a long working distance (>2 mm), high spatial resolution (1-2 μm across the field of view), high contrast (MTF10 < 1.5 μm), and a broad achromatic range (450-650 nm). Additional design parameters included an >0.4 numerical aperture, 5x magnification (corresponding to an ∼900 μm x 700 μm field of view), and 20 mm track length.

Our optical design included six custom microlenses to meet these goals (**Fig. 1a**; **Suppl. Table 1**). The lens surface profile, radius of curvature, distance between surfaces, and glass types were optimized using Zemax modeling. A #0 coverslip (∼100 μm thickness) was included in the design to minimize optical aberrations when imaging through live animal-implanted glass windows. Additionally, we accounted for a filter cube required for fluorescence imaging, placed in the collimated space between the lenses. Following optical modeling, the lenses and brass spacers were fabricated and assembled into two custom-made barrel holders minimized in size and weight (**Fig. 1b-c**). Each assembled barrel weighed <0.2 g.

**Fig. 1.**
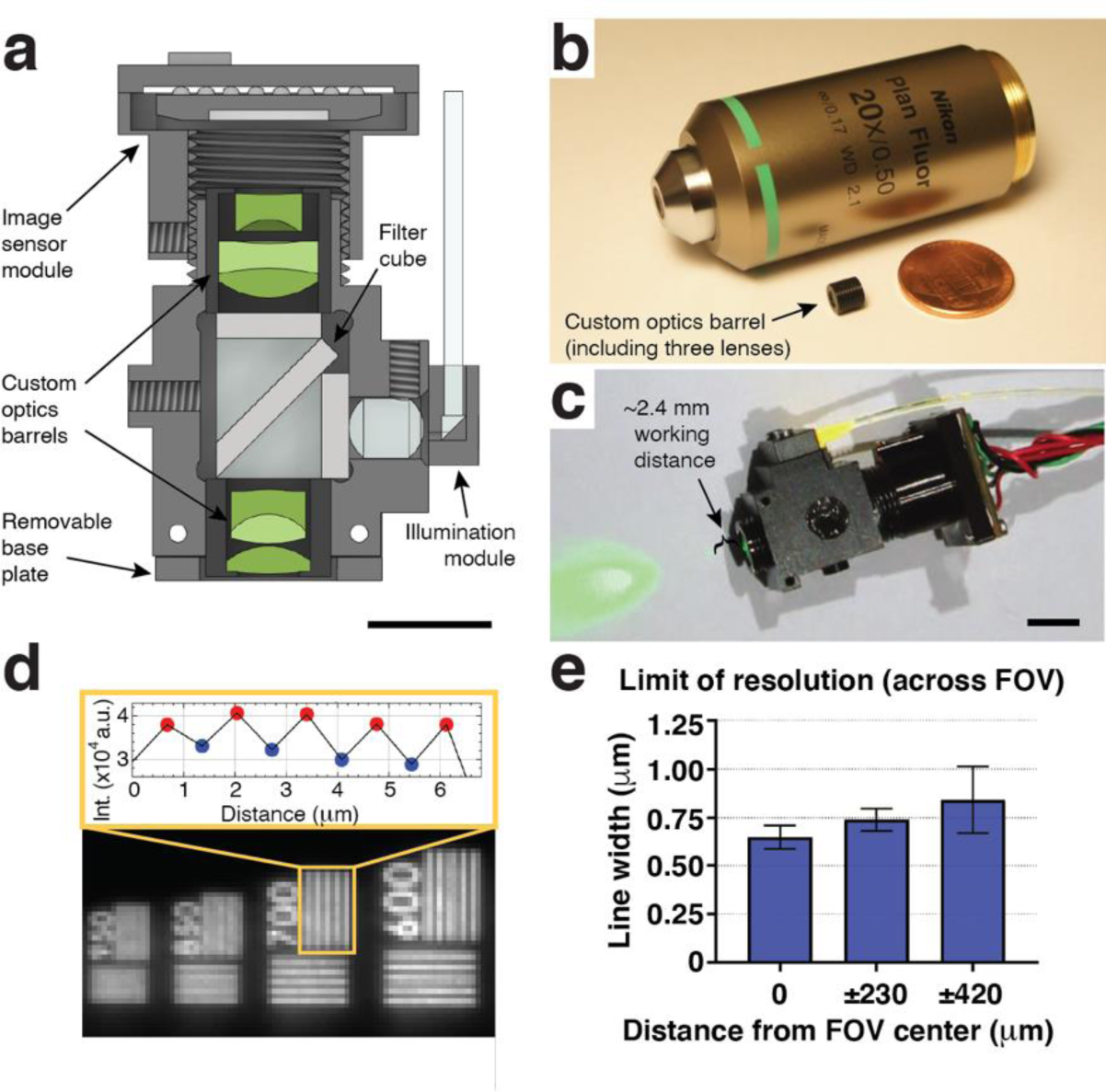
Wearable microscopes with long working distance custom-compound-microlenses for high-resolution, high-contrast measurements in freely behaving mice. **a**, Cross-section of the wearable microscope and its custom compound micro-optics (green) for high-resolution, multi-color, and long working distance imaging. Scale bar, 5 mm. **b**, Image of the <0.2-grams objective barrel including three miniature lenses next to a regular microscope objective and one-cent coin. **c**, Image of the fully assembled device with ∼2.4-mm working distance for high-resolution imaging through intermediary optics. Scale bar, 5 mm. **d**, *Bottom*, image detail of a high-resolution microscopy target demonstrating the wearable microscope’s resolving power. *Top*, intensity values of peaks and troughs across the indicated 700 lp/mm bar set. **e**, Limit of resolution across the field of view (FOV). Displayed values are averages across the horizontal and vertical line target results from comparable positions to the left and right of the FOV center. Spatial frequencies were converted to line widths. The data are presented as mean ± s.e.m.

Next, we characterized the performance of the custom microlens system. Based on our Zemax calculations, the predicted lateral and axial point spread function (PSF) in the center of the field of view (FOV) was 0.56 μm and 5.16 μm, respectively. Experimentally, we measured 0.68 ± 0.01 μm and 7.85 ± 0.33 μm (**Suppl. Fig. 1; Fig. 3**), indicating that lens fabrication and assembly only introduced minor errors. These measurements used a glass blank of similar thickness and composition in lieu of the filter cube, a separate potential source of aberrations. Our interchangeable filter cube consisted of multi-band excitation and emission filters, a custom multi-band dichroic beamsplitter, and two fused silica microprisms, joined together using an index-matching optical adhesive. This all-glass assembly minimized refractive index mismatch at the dichroic’s interfaces, preventing lateral beam offset and associated aberrations. A flexible PMMA fiber was used to deliver excitation light (**Fig. 1a, c**), providing flexibility in multi-color imaging applications. This design offers several advantages: a) It allows simultaneous coupling of multiple high-power light sources, such as DPSS lasers, enabling efficient excitation of co-expressed fluorophores (or opsins); and b) it provides flexibility in selecting functional reporters (and actuators).

To match the optical system’s high resolution, we chose a CMOS sensor with a 3.75 μm pixel spacing, corresponding to 0.69 μm at the object plane. This sensor, available in both RGB and monochrome versions, offered high-sensitivity (5.48 and 6.7 V/lux-sec for the RGB and monochrome version, respectively) and frame rates (45 fps at 1280 x 960-pixel resolution) and was mounted on a custom miniature printed circuit board (PCB) (**Methods**). Using a high-resolution microscopy target with spatial frequencies up to 3,300 lp/mm, we found that our integrated device had a limit of resolution (LOR) of 0.65 ± 0.07 μm and 0.84 ± 0.28 μm in the center and at the edges of the FOV, respectively (**Fig. 1d-e**). Using a regularly spaced grid target with 100 μm spacing, the FOV of our integrated system was 911 μm x 703 μm.

Next, to determine our microscope’s contrast limit, we measured the modulation transfer function (MTF) using the Slanted Edge test and MTF10 metric (i.e., contrast at 10%). Although MTF is frequently reported at 0%, in our experience, MTF values below 10% are affected by sensor noise limits and optical phase reversal. The MTF10 of our integrated device was 303 lp/mm tangential (1.65 μm line width) and 373 lp/mm sagittal (1.34 μm line width) in the center of the FOV (**Suppl. Fig. 2**).

Our microscope’s working distance (WD) was determined to be 2.365 mm (from the edge of the objective barrel to the microscopy target in air) using a precision differential actuator and grid target. This exceptionally long WD enables imaging through implantable micro-optics, such as microprisms (see below). Our fully assembled device measured 7 x 14 x 20 mm and weighed 2.96 grams, making it suitable for applications in mice. Together, these data demonstrate that our custom-compound-microlens system simultaneously provides exceptional spatial resolution, contrast, and working distance while retaining the large field of view, small form factor, and low weight compared to existing wearable microscopes.

### Multiplex imaging in behaving mice

Sensorimotor processing involves the dynamic interplay of numerous cell types’ activity. To evaluate our integrated system’s ability to perform multiplex measurements, we first imaged a monolayer of cell body-sized (15 μm) fluorescent micro-particles emitting three different spectral bands. As shown in **Fig. 2a-c**, the various colored microspheres could be brought into focus simultaneously, demonstrating the optical system’s low axial chromatic aberration across the FOV. To confirm this feature in a densely labeled environment, we imaged spinal tissue sections with neurons and astrocytes stained in two colors (**Fig. 2d**). The stained cells and their processes were readily distinguishable, further demonstrating our wearable device’s high resolution and contrast. As an additional test, we conducted multi-color imaging in behaving animals. GFAP-GCaMP6f mice, which constitutively express the green fluorescent calcium indicator GCaMP6f in astrocytes, were injected into the L3-L5 lumbar spinal cord with an AAV9-CaMKII-H2B-GCaMP7f-TagRFP vector, resulting in concurrent nuclear-localized GCaMP7f and TagRFP expression in excitatory neurons. Two to three weeks after the injection, we recorded noxious mechanical stimulus (tail pinch) evoked calcium activity in the superficial dorsal horn. To extract neurons’ and astrocytes’ evoked activity with minimal signal crosstalk (e.g., from out-of-focus cells), we defined computational exclusion criteria (e.g., TagRFP intensity level) to select appropriate regions of interest (ROIs) (**Fig. 2e-g**) (**Methods**). Consistent with our previous work^4, 9^, astrocytes showed widespread calcium activity to noxious tail pinch (p > 500 g) across the FOV, while running alone did not evoke concerted excitation in the superficial dorsal horn. In contrast, calcium transients in excitatory neuron nuclei were sparse^17^ and had lower signal amplitude. Their onset latencies and durations were comparable to astrocytes (**Fig. 2g**). Together, these data demonstrate that our custom-compound-microlens system and integrated microscopes allow simultaneous imaging of at least three colors, including calcium activity in different genetically defined cell populations of behaving mice, all with a single image sensor.

**Fig. 2.**
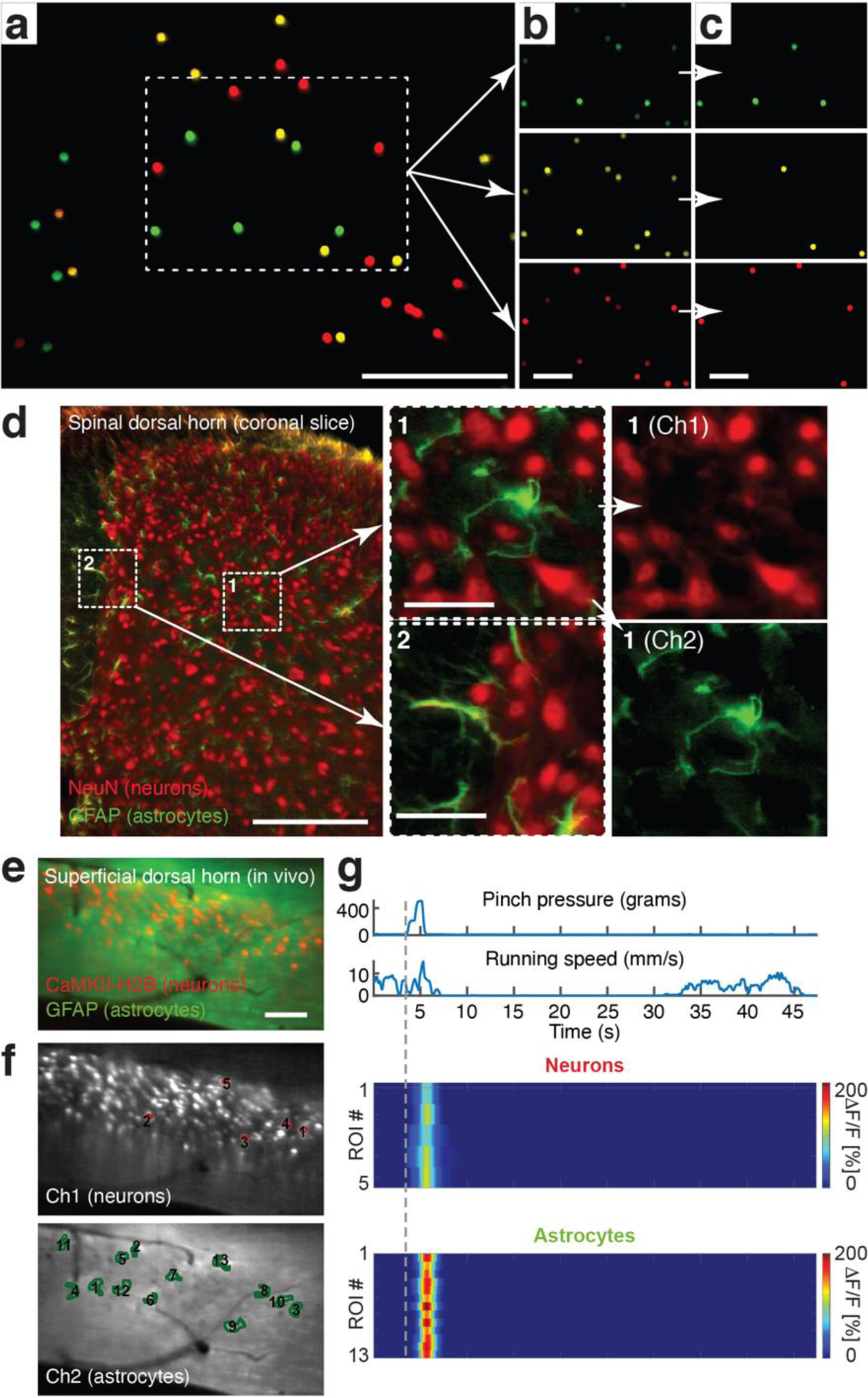
Wearable microscopes with custom-compound microlenses permit multi-color imaging with a single image sensor. **a**, Example fluorescence image of 15 μm-diameter polystyrene beads labeled with three distinct fluorophores demonstrating three-color imaging with a single RGB image sensor and dual-band filters. Scale bar, 250 μm. **b-c**, Individual color channels before (b) and after (c) color unmixing (**Methods**). Scale bars, 100 μm. **d**, *Left*, example fluorescence image of a 20-μm-thick mouse spinal cord section stained for astrocytes (GFAP; green) and neurons (NeuN; red) with two distinct fluorophores (Alexa Fluor 488 and 633) demonstrating two-color imaging *in vitro*. Scale bar, 250 μm. *Center*, blow-up of the two indicated subregions on the left. Scale bars, 50 μm. *Right*, color-separated images for subregion 1. **e**, Example fluorescence image from a time-lapse recording showing neuronal nuclei (red) and astrocytes (green) in the spinal dorsal horn of a behaving mouse demonstrating multiplex imaging *in vivo*. Imaging was performed about three weeks after AAV9-CaMKII-H2B-GCaMP7f-TagRFP injection into the lumbar spinal cord of GFAP-GCaMP6f mice. Scale bar, 100 μm. **f**, Color-separated images with neuronal (red) and astrocyte (green) regions of interest (ROIs) indicated. **g**, Noxious tail pinch-evoked neuronal nuclear and astrocyte calcium transients in the ROIs shown in f. Pressure stimuli were applied to the mouse’s proximal tail (**Methods**).

### Chronic translaminar imaging in the spinal cord of behaving mice through implanted microprisms

While wearable microscopes have uncovered sensory-evoked activity in the superficial dorsal horn of awake mice^4^, sensory activity in the spinal cord is not limited to these superficial regions. Measuring cellular activity in deeper and simultaneously across laminae at high speeds requires devices with sufficient WD to image through implanted micro-optics. To test our integrated system’s ability to enable such measurements, we first constructed tissue analogs in which reflective glass microprisms of different sizes were embedded (**Fig. 3a-c**) (**Methods**). These tissue phantoms also allowed us to quantify signal attenuation, imaging depth, and depth-dependent resolution in a scattering medium (**Fig. 3d-o**). Embedded 6 μm fluorescent beads were readily visible up to ∼350 μm distance from the vertical microprism face when imaging through a 0.7 mm x 0.7 mm x 0.7 mm (W x D x H) (**Fig. 3b, e, h, k, n**) or custom 0.7 mm x 0.7 mm x 1.7 mm microprism (**Fig. 3c, f, i, l, o**).

**Fig. 3.**
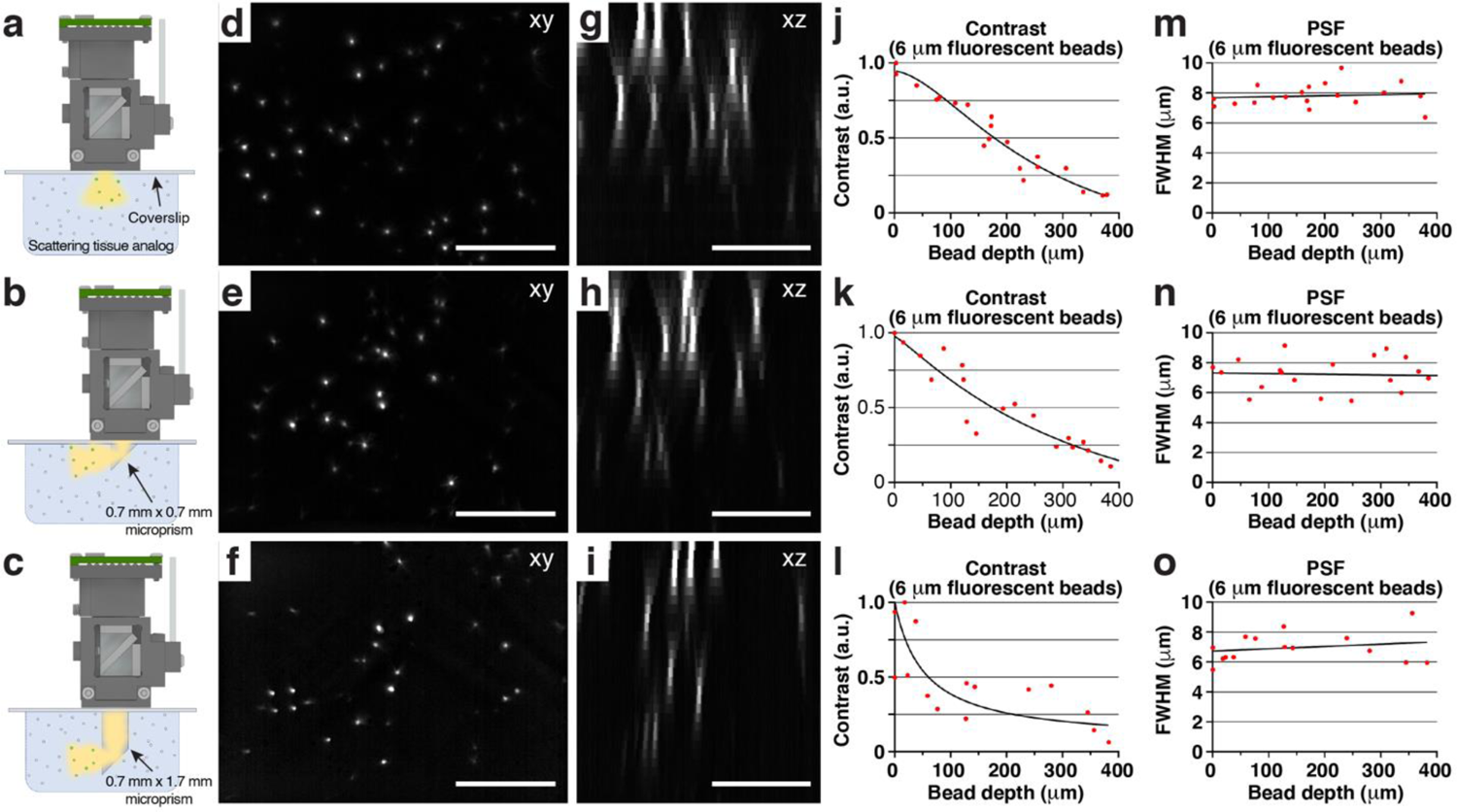
Wearable microscopes with custom-compound microlenses permit deep imaging through implanted microprisms. **a-c**, Schematics showing the experimental approach for characterizing and comparing three imaging conditions in scattering tissue phantoms (**Methods**): *Top*, imaging through a coverslip; *center*, imaging through a coverslip with an attached 0.7 mm x 0.7 mm x 0.7 mm microprism (W x D x H); *bottom*, imaging through a coverslip with an attached 0.7 mm x 0.7 mm x 1.7 mm microprism. **d-f**, example images of tissue phantom embedded 6 μm-diameter fluorescent beads. Each image is a maximum intensity projection through a z-stack acquired as shown in a-c by translating the microscope axially. Scale bar, 250 μm. **g-i**, maximum intensity side projections of the acquired z-stacks. Scale bar, 150 μm. **j-l**, bead contrast as a function of imaging depth. **m-o**, lateral FWHM of the 6 μm-diameter fluorescent beads as a function of imaging depth.

Next, we developed a surgical approach for microprism-assisted high-speed imaging across spinal laminae *in vivo*, including regions inaccessible by other high-resolution techniques. We found that inserting a 0.7 mm x 0.7 mm x 0.7 mm microprism at the lateral interface between the spinal gray and white matter was associated with minimal loss of NeuN-positive neurons two to four weeks after implantation (**Fig. 4a**; **Suppl. Fig. 3**). Microglia and astrocyte inflammatory responses were transient and confined to regions near the microprism-tissue interface, steadily declining over the four-week implantation period (**Suppl. Figs. 4-5**). Microprism-implanted mice did not show overt signs of sensory or motor deficits, as quantified by von Frey and kinematic weight-bearing tests (**Suppl. Fig. 6**). To determine whether our approach allows translaminar activity measurements in behaving animals, we performed optical recordings with our new wearable microscopes in the lumbar spinal cord of calcium indicator-expressing transgenic mice. Optical recordings were performed at 75 μm and 125 μm distance from the vertical prism face (**Fig. 4a-c**). Given these recording positions and the spinal cord’s known functional laminar arrangement, sensory stimuli and motor-evoked cellular activity would be expected to manifest in central and lower FOV regions, respectively, provided the investigated cells process corresponding sensorimotor information (**Fig. 4a-b**; **Suppl. Fig. 3**). To comprehensively quantify the spatial and temporal properties of cells’ activity patterns, we tiled the FOV with equally sized (10 μm x 10 μm) ROIs, excluding blood vessel regions (**Methods**). We chose noxious tail pinch (p > 500 g) as sensory stimuli for comparison with our previous work^4^. Starting with GFAP-GCaMP6f mice (N=5), we found that astrocytes exhibit calcium excitation across ∼200 μm-wide central FOV regions (**Fig. 4d-f**; **Suppl. Movie 1**), corresponding to sensory laminae. This areal activation was remarkably robust across animals, suggesting high reproducibility of our surgical preparation (**Fig. 4f**). Population transients evoked by noxious pinch had a 3.34 ± 0.21 s onset latency (**Fig. 4g**, left) and 2.79 ± 0.09 s duration (**Fig. 4h**, right), whereas individual ROI transients lasted 2.2 ± 0.02 s (**Fig. 4h**, left). In contrast, running-evoked astrocyte excitation in the same mice occurred in ∼300 μm-wide lower FOV regions (**Fig. 4i-k**; **Suppl. Movie 2**), likely corresponding to premotor areas. Average transients evoked by 2-4 s-long runs had 3.39 ± 0.19 s onset latency (**Fig. 4g**, right) and 2.75 ± 0.12 s duration (**Fig. 4l**, right), while individual ROI transients lasted 2.53 ± 0.03 s (**Fig. 4l**, left). Not every run triggered an astrocyte response, particularly after short inter-run rest periods (**Fig. 4j**), an effect previously described for motor-related brain areas^18, 19^. Next, we performed imaging in Tac1-GCaMP6f mice (N=3) with calcium indicator expression in Tac1-expressing neurons involved in pain processing^20, 21^. Noxious tail pinch (p > 500 g) evoked calcium excitation across ∼300 μm-wide central FOV regions (**Fig. 5a-c**; **Suppl. Movie 3**).

**Fig. 4.**
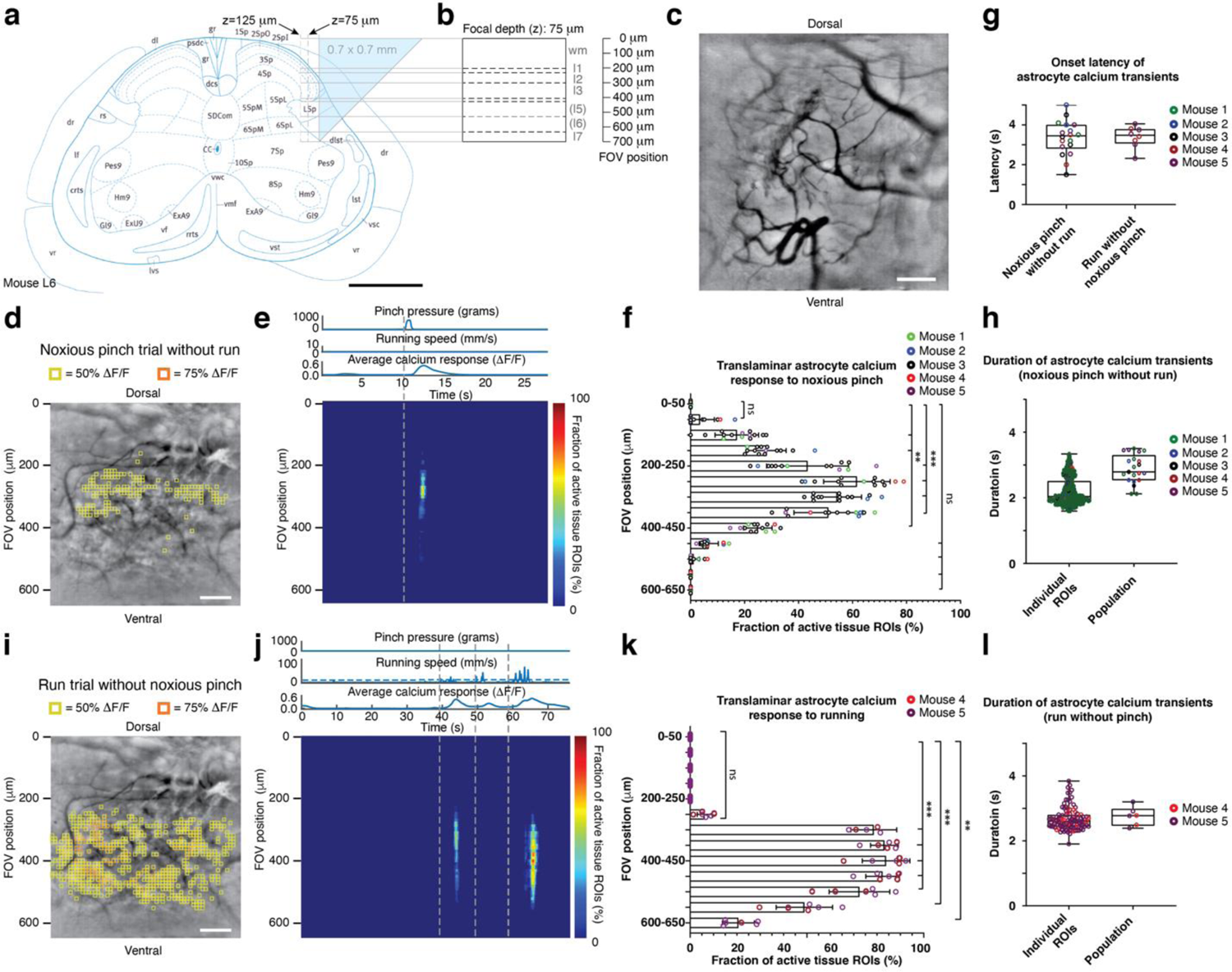
High-speed translaminar imaging reveals region-specific sensory and motor evoked activity in behaving GFAP-GCaMP6f mice. **a**, Schematic showing the translaminar imaging approach. An 0.7 mm x 0.7 mm x 0.7 mm glass reflective microprism was implanted at the lateral interface between the spinal gray and white matter at the L4-L5 spinal level (**Methods**). The microprism’s reflective hypotenuse tilts the microscope’s imaging plane by 90 degrees allowing high-speed measurements across laminae. Imaging was performed at 75 or 125 μm focal depth from the vertical microprism face. Scale bar, 0.5 mm. **b**, Predicted extent and borders of spinal laminae across the 700 μm field of view (FOV) for a 75 μm focal depth. **c**, Example fluorescence image showing the translaminar blood vessel pattern in a GFAP-GCaMP6f mouse at 75 μm focal depth four weeks after microprism implantation. Scale bar, 100 μm. **d**, Average intensity projection image from a different GFAP-GCaMP6f mouse at 75 μm focal depth overlaid with 10 μm x 10 μm ROIs. Only ROIs with at least 50% (yellow) or 75% ΔF/F (orange) in response to a noxious tail pinch (p > 500 g; duration: 1.5 s ± 0.5 s) are shown. In this example, the tail pinch did not evoke a locomotor response. Scale bar, 100 μm. **e**, Noxious tail-pinch evoked activity across tissue depth and time for the example recording shown in d. Each row depicts the percent of active ROIs (≥50% ΔF/F) for a given tissue depth. The corresponding pressure stimulus, locomotor activity, and average calcium transient across the FOV are shown above the activity heat map. Running speed was recorded by placing the animal on a spherical treadmill. **f**, Population data showing the average percent of active ROIs across tissue depths. Significant activity occurred in central FOV regions, corresponding to the upper spinal laminae. **g**, Population data of average calcium response onset latency for pinch-only and run-only trials. **h,** Population data showing individual ROI and average calcium transient duration for pinch-only trials. **i,** Activity map from the same animal as in d for a spontaneous run trial without pressure stimulus application. Same focal depth as in d. Scale bar, 100 μm. **j**, Running-evoked astrocyte activity across tissue depth and time. The pressure sensor readout, locomotor activity, and average calcium signal across the FOV are shown above the activity heat map. **k**, Population data showing the average percent of active ROIs across tissue depths for run trials. Significant activity occurred in FOV regions corresponding to deep spinal laminae involved in motor processing. **l**, Population data showing individual ROI and average calcium transient duration for run-only trials (2-4 s-long runs). The data in f, g (left), h is from 1,105 ROIs, 18 recordings, and 5 mice. The data in g (right), k, l are from 855 ROIs, 8 recordings, and 2 mice. All shown data were acquired four weeks after microprism implantation. Paired t-tests determined *P* values, and all bar plots are presented as mean ± s.e.m. The box and violin plots mark the median and the 25th and 75th percentiles, and the whiskers cover the minimum and maximum of the data.

**Fig. 5.**
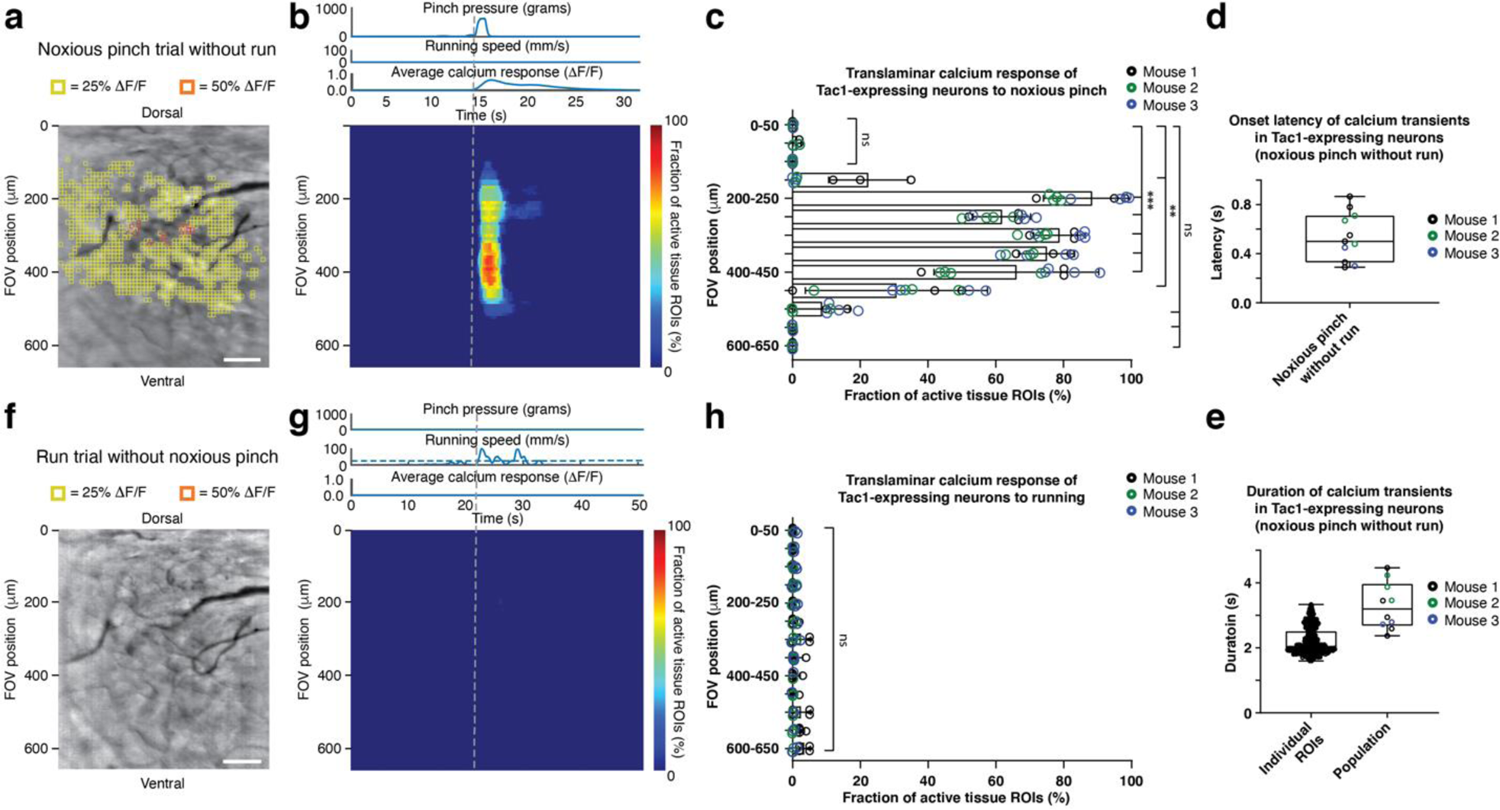
High-speed translaminar imaging reveals noxious mechanical stimulus-evoked activity in the spinal cord of behaving Tac1-GCaMP6f mice. **a**, Average intensity projection image from a Tac1-GCaMP6f mouse overlaid with 10 μm x 10 μm ROIs. Only ROIs with at least 25% (yellow) or 50% ΔF/F (orange) in response to a noxious tail pinch (p > 500 g; duration: 1.5 s ± 0.5 s) are shown. In this example, the tail pinch did not evoke a locomotor response. The focal depth was 75 μm. Scale bar, 100 μm. **b**, Noxious tail-pinch evoked activity across tissue depth and time for the example recording shown in a. Each row depicts the percent of active ROIs (≥25% ΔF/F) for a given tissue depth (Methods). The applied pressure stimulus, locomotor activity, and average calcium transient across the FOV are shown above the activity heat map. **c**, Population data showing the average percent of active ROIs across tissue depths. Significant activity occurred in FOV regions corresponding to the upper spinal laminae. **d**, Population data showing the average calcium transient onset latency for pinch-only trials. **e**, Population data showing individual and average calcium transient duration for pinch-only trials. **f**, Activity map from the same animal as in a for a spontaneous run trial without pressure stimulus application. Same focal depth as in a. Scale bar, 100 μm. **g**, Running-evoked activity across tissue depth and time. The pressure sensor readout, locomotor activity, and average calcium signal across the FOV are shown above the activity heat map. **h**, Population data showing the average percent of active ROIs across tissue depth for run-only trials. No significant calcium activity occurred within the FOV in response to running. The data in c, d, e are from 1,299 ROIs, 11 recordings, and 3 mice, while the data in h are from 8 recordings in 3 mice. All shown data were acquired four weeks after microprism implantation. Paired t-tests determined *P* values, and all bar plots are presented as mean ± s.e.m. The box and violin plots mark the median and the 25th and 75th percentiles, and the whiskers cover the minimum and maximum of the data.

Average transients across ROIs had an 0.54 ± 0.06 s onset latency (**Fig. 5d**) and 3.29 ± 0.23 s duration (**Fig. 5e**, right), whereas individual ROI transients lasted 2.20 ± 0.02 s (**Fig. 5e**, left), indicating signal propagation within and across laminae (**Fig. 5b**). Notably, unlike astrocytes, Tac1-expressing neurons did not show running evoked activity in premotor FOV areas (**Fig. 5f-h**).

Together, these data demonstrate that our approach provides real-time visualization of how different somatosensory programs activate different cell types in spatially restricted dorsoventral regions of behaving mice. Astrocytes showed acute mechanical pain and motor evoked calcium excitation in superficial and deep laminae, respectively. Tac1-expressing neurons responded to acute mechanical pain with upper but not deep lamina activation. These neurons also did not respond to locomotion, making them unlikely contributors to astrocytes’ motor evoked responses.

## Discussion

In this study, we have developed methods for high-speed translaminar activity measurements from genetically defined cell types in the spinal cord of behaving mice. Using chronically implanted micro-prisms with a 700 μm FOV, we could image sensory and motor evoked activity in sensory and premotor regions, respectively (**Figs. 4-5**). Our histological, behavioral, and functional data indicate that spinal microprism implantation is remarkably well tolerated by adult mice (**Suppl. Figs. 3-6**). To enable translaminar imaging in freely behaving animals, we developed and thoroughly characterized wearable microscopes with custom-compound microlenses (**Figs. 1-3**; **Suppl. Figs. 1-2**). This new integrated system offers a 2.4 mm working distance, ∼1.5 μm lateral resolution and MTF10 contrast, and 500-620 nm achromatic range, thereby overcoming multiple challenges of previous wearable microscopes. The combination of these innovations allowed us to uncover that dorsal horn astrocytes in behaving mice show distinct somatosensory program-dependent and lamina-specific calcium excitation (**Fig. 4**).

Additionally, we found that Tac1-expressing neurons exhibit upper laminae-restricted activity to acute mechanical pain but not locomotion (**Fig. 5**). Most previous wearable one-photon microscopes, including our own^4, 22, 23^, rely on gradient-index lenses^9, 14, 15^. While the optical capabilities of these lenses have seen significant progress, performance limitations remain, such as their narrow achromatic range, short working distance, or variable resolution and contrast across the field of view^16^. Wearable one-photon microscopes with more traditional lens systems have emerged but tend to use few or off-the-shelf optical elements^9, 14, 15^, likely because of the difficulty of modeling, optimizing, or producing more complex compound-microlens systems, thereby constraining the number of desirable optical features that can be achieved concurrently. We, therefore, took an *ab initio* approach that involved paraxial calculations and Zemax modeling (**Methods**). Our custom-compound-microlens system provides a wide achromatic range (from the visible to near-infrared), high spatial resolution and contrast, and an exceptionally long working distance (**Figs. 1-3**; **Suppl.** Figs. 1-2).

Partly because of these technical limitations, previous imaging approaches could not record cellular activity across sensory and premotor areas, preventing vital biological questions from being addressed^9^. For example, recent gene expression and electrophysiological studies have indicated that astrocytes show lamina-specific properties^7, 24, 25^, raising questions about how this heterogeneity might be linked to differences in functional signaling. Our data show that acute pain and animal locomotor activity evoke astrocyte calcium excitation in distinct dorsal horn regions (**Fig. 4**), supporting astrocytes’ responsiveness and functional adaptation to region-specific neural circuits and signaling. Likewise, genetic, immunohistological, and electrophysiological approaches have implicated Tac1-expressing neurons in pain signaling^20, 21^. Our data show that noxious mechanical stimuli lead to rapid and widespread calcium activity across dorsal horn laminae in these neurons (**Fig. 5**), the first such measurements in behaving animals.

High-speed translaminar imaging was achieved by microprism implantation. Developing this approach for the spinal cord was a non-trivial endeavor given its small cross-sectional dimensions compared to the brain, the limited space within mouse vertebrae, and region-dependent tissue displacements during animal behavior^4, 26^. We found that 0.7 mm x 0.7 mm x 0.7 mm microprisms provide an acceptable FOV while minimizing tissue disruption. No significant loss of neuronal cell bodies was observed 2-4 weeks after implanting microprisms at the lateral interface between the spinal gray and white matter (**Suppl. Fig. 3**). Glial reactivity was transient and confined to regions near the tissue-microprism interface (**Suppl. Figs. 4-5**). Sensory and motor tests before and after microprism implantation also revealed no overt behavioral deficits (**Suppl. Fig. 6**), and our calcium imaging data suggest that spinal neurons and astrocytes remain responsive to sensory input and motor actions (**Figs. 4-5**). Astrocyte response properties to noxious tail pinch (e.g., onset latency, duration) were comparable to data from non-implanted mice^4^. Nevertheless, microprism implantation causes some circuit disruption (e.g., to white matter tracts near the implantation site) and should be considered in future biological applications on a case-by-case basis, as is generally advisable for any imaging approach that uses implanted micro-optics.

Repeated imaging through spinal cord-implanted microprism is feasible over at least four weeks. This recording period may enable the study of prolonged structural and functional biological processes, such as disease- or treatment-related dynamics (e.g., amyotrophic lateral or multiple sclerosis progression, transplanted neural stem cell integration)^9, 10, 27^. Optical recordings from the spinal ventral horn might be feasible with custom micro-optics. As demonstrated in **Fig. 3**, our wearable microscopes have sufficiently long WD to image through 0.7 mm microprisms with a 1 mm vertical extension, potentially allowing the study of motor neuron activity in relation to limb movement. Our surgical and optical methods may also apply to thoracic and cervical spinal cord regions, allowing the study of sensorimotor activity related to internal organ or forelimb function. However, this will likely require the development of imaging chambers specifically adapted to the different vertebra anatomy in these regions. Furthermore, given the rapidly expanding number, color palette, and signal-to-noise ratio of genetically encoded indicators (e.g., for neurotransmitters and neuromodulators)^28^, our wearable microscopes’ multiplex imaging capability might in the future enable the study of input-output relationships (e.g., integration of molecular signals by a given cell type or modulation of local cellular activity by descending projection fibers from the brain). In summary, our novel surgical and optical techniques should enable a wide range of biological interrogations, providing unprecedented opportunities for elucidating cellular and molecular mechanisms of sensorimotor information processing in the spinal cord of behaving animals.

## Methods

### Miniature microscope design

Our wearable microscopes include six major components: the custom optics barrels (including six microlenses), fluorescence filter cube, main body, image sensor module, illumination module, and base plate (**Fig. 1a**).

#### Optical design

The optical design was performed using Zemax optical modeling software and manual calculations. During the design phase, geometric distortion was deliberately allowed to increase to 5.7% to gain improvement across the other Seidel aberrations (spherical, coma, astigmatism, and field curvature). Our final optical design utilized six lens elements, each from a different type of glass and housed in two separate barrels (Optics Technology Inc.).

The filter cube consisted of a custom 4.2 x 6.84 x 1 mm dual-band dichroic beamsplitter (59012-custom, Chroma Technology Corp.), 4.1 x 4.2 x 1.1/1.0 mm dual-band fluorescence excitation and emission filters (59012x and 59012m, Chroma Technology Corp.), and two uncoated 4.2 x 4.2 x 4.2 mm fused silica prisms (Tower Optical Corp.). This all-glass filter cube configuration was designed to correct the lateral beam path offset and optical aberrations introduced by the dichroic beamsplitter. Dichroic beamsplitters are typically designed for 45-degree incidence from air. We, therefore, designed a custom coating that matched the 59012bs transmission spectrum allowing the filter to be used for a 45-degree incidence from glass. Filter cube assembly was done manually using index-matched UV-cure adhesive (NOA146H, Norland Products Inc.) to minimize internal reflections. The outside of the filter cube was painted with a custom black paint (black pigment dispersed in NOA146H, Norland Products Inc.) to absorb stray light reflections. A retainer ring was introduced to reduce filter cube-related optical aberrations (e.g., color shifts) at the edges of the FOV. This ring decreased the effective NA from 0.46 to 0.41, a ∼21% reduction in the light collection.

The illumination module was designed to hold a hand-polished poly(methyl methacrylate)-core (PMMA-core) multi-mode fiber (MMF) with 735 μm core and 0.51 NA (Eska, Mitsubishi International PolymerTrade Corp.). We selected a PMMA-instead of a silica-core MMF because its flexibility aided the animal‟s mobility. The large core size and NA provided efficient light coupling (e.g., from LEDs). The illumination module also included a 1.4 mm 45°-fold-mirror, constructed from two 45°-45°-90° 1 x 1 x 1 mm micro-prisms with a reflective-coated hypotenuse (Tower Optical Corp.), mounted inside the module. The fold mirror was coupled to the optical fiber using index-matched UV-cure adhesive (NOA81, Norland Products Inc.), while the collector lens (45-664, Edmund Optics) was secured inside the microscope’s main body. The fold mirror allowed the MMF to be vertically coupled to the horizontal collector lens, an intentional design to also aid the animal‟s mobility.

As the illumination sources, we either used two high-intensity table-top LEDs or lasers. The LED light source consisted of blue (470nm) and yellow (554nm) mounted LED packages (M470L4 and MINTL5, Thorlabs) combined in free-space and coupled into a silica-core intermediary MMF patch-cord (M59L01, ThorLabs). The patch-cord was then mated to the illumination module‟s MMF using an SMA to SMA mating sleeve (ADASMA, ThorLabs). The purpose of the intermediary MMF patch-cord was to allow for the precise alignment of the LEDs to be unaffected when switching light sources (e.g., between the LEDs and lasers). A 240 μm core MMF (Eska, Mitsubishi International PolymerTrade Corp.) was used with table-top lasers (473 nm and 556 nm DPSS lasers; MBL-FN-473 and MGL-FN-556, respectively; CNI Optoelectronics Tech. Co.).

#### Optomechanical design

The housing included four modules (main body, sensor mount, illumination module, and base plate) and was designed in CAD software (Autodesk Inventor). This modular design simplifies the optimization process and allows for future upgrades without the need to develop an entirely new device. The main body holds the optics barrels, the filter cube, and the illumination collector lens. It was designed with an open central compartment to access the interchangeable fluorescence filter cube easily. The main body also includes features for mating with the sensor mount, base plate, and illumination modules. The sensor mount holds the image sensor and blocks stray light. It contains threading and a locking setscrew to allow fine focus adjustments after mounting. The illumination module was designed to provide flexibility in using LED or laser light sources in multiplex imaging applications. The detachable base plate allows reproducible mounting, facilitating chronic imaging experiments. For ease of fabrication, three-axis computer numerical control (CNC) milling was selected. This technique influenced the design process as the complex geometry needed to be confined to dimensions accessible to the CNC mill. Most housing components were fabricated from the lightweight polyether ether ketone (PEEK) (Zera Development Company). The sensor mount was fabricated from aluminum to aid in dissipating the heat generated by the image sensor.

#### Electronic design

We selected a CMOS sensor (MT9M024, Aptina Imaging) that matches our optical resolution (3.75 um pixel spacing, corresponding to 0.69 μm at the object plane; 1280 x 960 pixels) and offers high sensitivity (5.48 and 6.7 V/lux-sec for the RGB and monochrome version, respectively) and frame rate (45 fps at full resolution). The sensor was mounted on a custom miniature printed circuit board (PCB), interfaced with an Aptina data-acquisition board (AGB1N0CS-GEVK), and controlled using the Aptina DevWare Software Package. For high-speed multi-color imaging (at 45 fps), we used an RGB version of the sensor. For high-sensitivity imaging through microprisms, we used its monochrome version. All data were recorded with 12-bit depth.

#### Microscope characterization

Optical system performance was characterized before any software-based post-processing (e.g., spatial down-sampling, PCA, or ICA). Image resolution was quantified via the point spread function (PSF) and the limit of resolution (LOR) test (**Fig. 1d-e**; **Fig. 3d-i, m-o**; **Suppl. Fig. 1**). To measure the imaging optics‟ PSF, we designed a test rig that secured the optics barrels in place. It also contained a simulant filter block (i.e., a glass blank of the same thickness and composition as the fluorescence filter block) and spacer on the object side to ensure a working distance of ∼2 mm to the target. A Zeiss LSM 780 confocal microscope was used to image sub-resolution (0.5 μm) fluorescent beads through the optics barrels. To measure the integrated system’s PSF, we generated a point source of light using a 0.5 μm pinhole (TC-RT01, Technologie Manufaktur), green fluorescence reference slide (2273, Ted Pella), and blue LED excitation light source (M470L3, Thorlabs). We then took images in evenly spaced axial intervals around the focus point of the assembled microscope. PSF values were calculated from lateral and axial maximum intensity image stack projections and are specified as the full width at half maximum (FWHM) (**Fig. 3d-i, m-o**; **Suppl. Fig. 1**).

To measure the LOR at the center and different locations across the FOV (210 ± 20 μm spacing), we recorded images of a high-resolution microscopy USAF target (TC-RT01, Technologie Manufaktur) with a maximum spatial frequency of 3300 lp/mm (**Fig. 1d-e**). The focus was set at the center and kept stable across the radial positions to account for field curvature and astigmatism. We plotted the line profile across the bar variance at each recording location and averaged along the bar length. We then averaged the peaks and troughs and calculated the contrast percentage using the Michelson contrast equation. This data analysis utilized the “Find Peaks” ImageJ plugin. We defined the LOR limit (highest spatial frequency) by the bar set where at least 5% contrast can be observed.

The contrast was quantified via the modulation transfer function (MTF) (**Suppl. Fig. 2**). Contrast is the object’s brightness relative to the background calculated via the Michelson contrast equation. To measure the MTF, we employed the Slanted Edge test using a razor blade, green fluorescence reference slide (2273, Ted Pella), and blue LED excitation light source (M470L3, Thorlabs). The focus was set at the center of the FOV and kept stable across different radial positions (the same positions as in the LOR test). Corresponding data were analyzed using the “Slanted Edge MTF” ImageJ plugin. We calculated both the sagittal and tangential MTF. While MTF 0% is frequently reported, MTF values below 10% are affected by sensor noise limits and optical phase reversal. MTF spatial frequency is therefore reported at 10%, mimicking biological samples of a specific labeling density. Corresponding line widths are equal to one-half the spatial period.

The FOV was measured by imaging a grid target, printed directly on the surface of the microscope slide without glass coverslip, and using a precision actuator (DRV3, Thorlabs). To demonstrate the multi-color imaging capability of our integrated device, we imaged cell body-sized (15 μm) fluorescent polystyrene microspheres of different colors (yellow-green F8844, yellow F21011, and red F8842 beads with 505 nm/515 nm, 515 nm/534 nm, 580 nm/605 nm excitation/emission peaks; ThermoFisher) (**Fig. 2a-c**). Corresponding data were processed in DevWare using the demosaicing algorithm “Anisotropic Diffusion.”

The working distance (i.e., the distance from the edge of the lower optics barrel to the object plane) was measured by placing the microscope in contact with a grid target (R1L3S3P, Thorlabs) and then translating it upwards with a precision differential actuator (DRV3, Thorlabs) until the image was in focus. The working distance depends on the position of the image sensor along its focusing range and the thickness of the cover glass. We used a #0 glass coverslip (∼100 μm thickness) for our measurements and positioned the image sensor in the middle of its focusing track.

The weight of the integrated system was determined by weighing the assembled device, including the custom optics barrels, filters, illumination module, housing, image sensor, and sensor PCB. We did not include the sensor wires or optical fiber in the measurement because external mounts or a commutator typically support the weight of these tether components.

### Experimental model and subject details

All procedures were performed following the National Institutes of Health (NIH) guidelines and were approved by the Institutional Animal Care and Use Committee (IACUC) at the Salk Institute. Mouse strains used in this study included GFAP-Cre (RRID: IMSR_JAX:012886), Tac1-Cre (RRID: IMSR_JAX:021877), and Ai95(RCL-GCaMP6f)-D mice (RRID: IMSR_JAX:024105)^29–31^. Mice were group-housed, provided with bedding and nesting material, and maintained on a 12-h light-dark cycle in a temperature (22 ± 1°C) and humidity controlled (45-65%) environment. All the imaging and behavioral experiments involved 6-11 weeks-old heterozygous male and female mice. Experimental mice used in individual experiments typically originated from different litters. Mice had marks for unique identification. No criteria were applied to allocate mice to experimental groups.

### Molecular cloning, AAV production, and titering

The cDNA for H2B-GCaMP7f-TagRFP was PCR amplified and subcloned into an AAV transfer vector downstream of the CaMKIIa promoter and upstream of WPRE and hGHpA sequences. This vector was co-transfected into HEK293-AAV cells (Vector Biolabs) along with a pAdeno-helper vector and a pRC-AAV9 rep-cap plasmid. Recombinant AAV9 production was then carried out using a protocol developed by the Byungkook Lim laboratory at UCSD. The recombinant AAV9-CaMKIIa-H2B-GCaMP7f-TagRFP-WPRE-hGHpA was titered by qPCR using primers designed to the hGHpA sequence. The titer of the virus was 4.9E+12 GC/ml.

### Stereotactic injections

For dual-color *in vivo* imaging (**Fig. 2e-g**), the AAV9-CaMKII-H2B-GCaMP7f-TagRFP vector was injected into the L3-L5 spinal cord (coordinates: ML 0.3, DV 0.15-0.3 mm; volume: 1 μl; dilution: 1:25). Surgical procedures closely followed previously established protocols^4^. Briefly, thin-wall glass pipettes were pulled on a Sutter Flaming/Brown micropipette puller (model P-97). Pipette tips were cut at an acute angle under 10x magnification using sterile techniques. Tip diameters were typically 15–20 μm. Pipettes that did not result in sharp bevels or had larger tip diameters were discarded. Millimeter tick marks were made on each pulled needle to measure the virus volume injected into the spinal cord. Mice were anesthetized with isoflurane (4-5% for induction; 1%-1.5% for maintenance) and positioned in a computer-assisted stereotactic system with digital coordinate readout and atlas targeting (Leica Angle Two). Body temperature was maintained at 36–37°C with a DC temperature controller, and ophthalmic ointment was used to prevent eyes from drying. A small amount of depilator cream (Nair) was used to remove hair at and around the incision site. The skin was then cleaned and sterilized with a two-stage scrub of betadine and 70% ethanol, repeated three times. Surgical scissors were used to make a small (around 10 mm) incision along the dorsal midline. Fascia connecting the skin to the underlying muscle was removed with forceps. The skin was held back by retractors. Using blunt dissection, lateral edges of the spinal column were isolated from connective tissue and muscle. Tissue from the vertebra of interest and one vertebra rostral and caudal to the site of spinal cord exposure was removed with forceps. The spine was then stabilized using Cunningham vertebral clamps, and any remaining connective tissue on top of the exposed vertebrae was removed with a spatula. An approximately 0.3 mm opening was made in the tissue overlying the designated injection site (e.g., using sterile Dumont #3 forceps). For injection, a drop of the virus was carefully pipetted onto parafilm (2–3 μl) for filling the pulled injection needle with the desired volume. Once loaded with sufficient volume, the injection needle was slowly lowered into the spinal cord until the target depth was reached. Manual pressure was applied using a 30-ml syringe connected by shrink tubing to slowly inject the viral solution over 5–10 min. The syringe‟s pressure valve was then locked. The position was maintained for approximately 10 min to allow the virus to spread and to avoid backflow upon needle retraction. Following the injection, spinal cord clamps were removed, muscle approximated, and the skin sutured along the incision. Mice were given subcutaneous Buprenex SR (0.5 mg/kg) and allowed to recover before placement in their home cage.

### Live animal preparation

Animals were implanted with a spinal and head plate under general anesthesia approximately one week before laminectomy, as previously described^4^. Buprenex SR (0.5 mg/kg) was given to minimize post-operative pain.

#### Multi-color imaging of superficial laminae

A laminectomy (2 wide x 4 mm long) was performed at the T12-T13 vertebra level, corresponding to spinal segments L3-L5^26^, four days before the initial recording session. The dura mater overlying the spinal cord was kept intact, and a custom-cut #0 coverslip was used to seal the laminectomy creating an optical window for imaging. The coverslip was replaced immediately before recording sessions to maximize optical clarity.

#### Translaminar imaging through implanted glass reflective microprisms

At least one day before surgery, a 0.7 x 0.7 x 0.7 mm glass microprism with aluminum-coated hypotenuse (cat. no. 4531-0021; Tower Optical) was UV-cured (NOA 81; cat. no. 8106; Norland Products Inc.) to a custom-cut #0 coverslip matching the intended laminectomy size (3 mm wide x 4 mm long). To minimize bubbles at the microprism-coverslip interface, the microprism was moved in a circular motion and then positioned at the desired location before UV curing. Excess glue around the microprism was removed with sterile absorbent paper points (cat. no. 50-930-669; Fisher Scientific). After UV curing, the microprism-coverslip assembly was placed in a sterile glass petri dish with a cover and heated on a hot plate at 50°C to allow further curing for 8-12 h. Following heat curing, the microprism-coverslip assembly was stored in a 70% EtOH solution. To implant this assembly, mice were anesthetized with isoflurane (4-5% for induction and 1-2% for maintenance in 100% oxygen) and placed in a stereotactic setup. Body temperature was maintained at 36-37°C with a DC temperature controller. A laminectomy (3 mm wide x 4 mm long) was performed at the T13-L1 vertebra level, corresponding to spinal segments L4-L6^26^.

Using a dissecting knife (cat. no. 10055-12; Fine Science Tools) attached to a stereotactic arm, a small incision was made 0.7 mm lateral to the central vein‟s center, coinciding roughly with the interface between the dorsal root ganglia (DRG) and spinal white matter in 6-8 weeks-old mice. The incision extended 0.7 mm in the rostrocaudal direction and 0.7 mm in depth, matching the microprism dimensions. Cerebrospinal fluid but not blood briefly exuded from the incision site upon retraction of the dissecting knife. The surgical site was washed several times with cold, sterile saline. The microprism implant, held by an 18G x 1 ½” blunt angled syringe needle (cat. no. 305180; Thermo Fisher Scientific) attached to a vacuum suction line, was positioned above the incision site. The implant was slowly lowered until fully inserted (∼0.7 mm depth). Excess fluid was removed using sterile absorbent paper points (cat. no. 50-930-669; Fisher Scientific). Once affixed to the surrounding bone with instant adhesive (cat. no. 3EHP2; Grainger), the suction holding the implant was turned off.

### Fluorescence imaging

#### *In vivo* imaging through a dorsal optical window

Data in behaving mice were acquired following previously established protocols^4^. Fiber-coupled blue (470nm) and yellow (554nm) LEDs (M470L4 and MINTL5, Thorlabs) were used for GCaMP7f and TagRFP excitation (**Fig. 2e-g**). Approximately 10-20 recordings were taken per imaging session, with each recording lasting around 1-2 min. The typical average power for a given recording and light source was <125 μW mm^-2^. No signs of phototoxicity, such as a gradual increase in baseline fluorescence, lasting changes in activity rate, or blebbing of labeled cells, were apparent in our recordings.

#### *In vivo* imaging through implanted microprisms

Data in behaving mice were acquired up to four weeks after microprism implantation (**Figs. 4-5**). The miniature microscopes‟ FOV allowed imaging of the entire microprism face. Recordings were taken at 75 μm and 125 μm focal depth from the vertical microprism-tissue interface, where 0 µm was defined as the point when tissue blood vessels first came into focus. A fiber-coupled 473 nm DPSS laser was used for GCaMP6f excitation. Each imaging session included approximately 10-20 recordings, with each recording lasting around 1-2 min. The typical average light power at the tissue surface was between 275-325 μW mm^-2^. All *in vivo* data was acquired at the image sensor‟s full resolution (1,280 x 960 pixels) and maximum frame rate (∼45 Hz).

#### Confocal imaging of stained tissue sections

Three-channel, 3 x 3 tiled z-stacks (15 images at 1 μm axial spacing) were acquired with a Zeiss LSM 710 confocal microscope to produce images of whole spinal cord sections (laser lines: 405 nm, 488 nm, 633 nm). Each image within the z-stacks had a 1,024 x 1,024-pixel resolution and was acquired using an Olympus 20x 0.8 NA air-matched objective.

### Sensory stimuli and behavioral tests

#### Sensory stimuli during *in vivo* imaging

Each imaging session consisted of up to 20 recordings. In a subset of recordings, mechanical stimuli were delivered to the animal‟s tail using a rodent pincher system (cat. no. 2450; IITC Life Science, Inc.). Pinch pressures were applied in the dorsoventral direction at approximately 6 mm from the base of the animal’s tail. Each pinch stimulus (typically one per recording) lasted around 1-2 s. Subsequent stimuli were delivered at least 1.5-2 minutes apart to minimize response adaptation. The order in which stimuli of different amplitudes were delivered was randomized. **Figs. 2e-g**, **4-5** show data from p > 500 mechanical stimuli.

#### Sensory tests before and after microprism implantation

To quantify the effects of microprism implantation on the animal’s sensory performance, we used the von Frey assay (**Suppl. Fig. 6a-b**). The Simplified Up-Down (SUDO) method^32^ was performed using Touch Test Sensory Evaluators (North Coast), starting on the sixth (0.4 g) fiber. The SUDO Method utilizes five stimulations per test, ensuring each animal undergoes the same number of hind paw stimulations. Animals were habituated to the behavioral testing room for at least 15 minutes in their home cage, then to the von Frey apparatus for at least 45 minutes with the researcher outside of the room, and the researcher’s presence for an additional 15 minutes. Stimulations were performed every 5 minutes, alternating between hind paws. Four female GFAP- and Tac1-GCaMP6f mice were tested before and 7 days after spinal plate implantation and again at two and four weeks after microprism implantation. Data from the two-week time point are an average of trials performed on days 13 and 15, whereas data from the four-week time point are an average of trials conducted on days 27 and 29 after microprism implantation. Animals were tested twice to decrease the effect of inter-day noise, with a rest day between trials. *In vivo* microscopy was performed on days 14 and 28. The researcher was blinded to the microprism implantation side (left or right side of the spinal cord).

#### Motor tests before and after microprism implantation

To quantify the effects of microprism implantation on the animal’s locomotor performance, we used the kinematic weight-bearing test (Bioseb) (**Suppl. Fig. 6c-f**). Animals were acclimated to the testing environment by placing their home cage in the testing room approximately 30 minutes before the recordings. The animals were then transferred to the behavior setup, allowing them to explore the new environment for about 15 minutes freely. Following this period, five runs were collected per animal and for five animals total. Runs were counted only if the animal moved from left to right in the setup without stopping. Stride length (cm), peak force (cN), and running speed (cm/s) were analyzed. Runs of a given animal were averaged. The same animals were measured before and after spinal plate implantation, and two and four weeks after microprism implantation.

### Immunohistochemistry

Two or four weeks after microprism implantation, mice were euthanized in their home cage following American Veterinary Medical Association (AVMA) guidelines. Transcardial perfusion was performed using 10% sucrose followed by 4% paraformaldehyde. L1-L6 spinal cord tissue was carefully extracted and incubated in 4% PFA overnight at 4°C. The tissue was then washed on a shaker with 1x PBS three times with 15 min per cycle, followed by incubation in 30% sucrose in PBS overnight at 4°C. Next, the tissue was flash-frozen in an OCT tissue-freezing medium, cryosectioned at 20 µm thickness, air-dried overnight, and then processed for immunostaining. Tissue sections were incubated overnight at 4°C with primary antibody diluted in blocking buffer, then washed in PBS containing 0.1% Tween-20, and incubated for two hours at 22-24°C in the dark with fluorophore-coupled secondary antibodies. Primary antibodies included NeuN (1:100; Novus Biologicals Inc.; cat. no. NBP1-92693; RRID: AB_11036146), GFAP (1:250; Thermo Fisher Scientific; cat. no. 13-0300; RRID: AB_2532994), and Iba-1 (1:200; FUJIFILM Wako Shibayagi; cat. no. 019-19741; RRID: AB_839504). Secondary antibodies included Alexa Fluor 633 goat anti-mouse (1:100; Thermo Fisher Scientific; cat. no. A-21052; RRID: AB_2535719), Alexa Fluor 488 goat anti-rat (1:100; Thermo Fisher Scientific; cat. no. A-11006; RRID: AB_2534074), and Alexa Fluor 405 goat-anti-rabbit (1:100; Thermo Fisher Scientific; cat. no. A-31556; RRID: AB_221605).

### Image data processing and analysis

#### *In vivo* imaging data

We used a custom Matlab script to convert the image sensor’s raw files from 12- to 16-bit, demosaic corresponding images (for RGB image sensor data only), and adjust their pixel range. The data were then further processed in Fiji. We converted the image data to TIFF, cropped the time-lapse recordings to the LED-/laser-on period and fluorescently labeled central areas of the FOV, followed by illumination correction, background subtraction, and color separation (for RGB image sensor data only). Full-frame image motion was reduced using Moco^33^. For multi-color image data, one channel was selected to determine and correct x-y image displacements across all channels. Image edge artifacts introduced by motion correction were cropped. Within-frame distortions were typically small and therefore not corrected. Motion-corrected calcium imaging data were analyzed using custom ImageJ and MATLAB software. Manually drawn cell body-sized ROIs were used to extract activity traces from neurons and astrocytes in dual-color recordings (**Fig. 2e-g**). We applied several quantitative exclusion criteria to ensure that the activity traces derived from these ROIs are from a defined cell type. Activity traces were considered from neurons if the average TagRFP fluorescence within respective ROIs exceeded the average TagRFP fluorescence across the FOV by at least 2 s.d. This retained only AAV9-CaMKII-H2B-GCaMP7f-TagRFP transduced cells with sharp boundaries (i.e., in-focus neurons). Activity traces were considered from astrocytes if the average TagRFP fluorescence within the respective ROIs was below 0.5 s.d.

To quantify calcium activity more comprehensively across the FOV, we used an unbiased tiling approach, similar to our previous work^34^ (**Figs. 4-5**). Each tile corresponded to a 10 x 10 μm ROI. We defined three ROI classes: over, near, and distant from blood vessels. First, we calculated the mean and s.d. of the recording’s average intensity image. Next, we determined individual ROIs’ average intensity values (A). If A < mean - s.d., it was considered a blood vessel-ROI. If A ≥ mean + (1+x)*s.d. (with x ∈ [0.3-0.7] depending on data set (e.g., blood vessel size/pattern)), it was classified as distant from blood vessels. ROIs with in-between values were considered near-blood vessel-ROIs. ROIs over and near blood vessels were excluded from data analysis. For all ROIs distant from blood vessels, we calculated their average fluorescence intensity over time. Corresponding activity traces were temporally smoothed using an 0.4 s sliding average. Local maxima within the traces were identified using Matlab’s “findpeaks” function. The fluorescence baseline and noise level (in s.d.) of a given trace were calculated using a 2 s period before stimulus or run onset. Local maxima were considered evoked activity if fluorescence intensity values surrounding the peak were at least 6 s.d. above baseline for ≥2 s. Calcium transient onset was defined as the point at which the fluorescence intensity trace immediately before the peak crossed a 2 s.d. threshold above baseline. Calcium transient duration was defined as the trace’s full width at half maximum (FWHM). Traces that showed a) activity within 2 s after recording onset (i.e., before stimulus or run onset), b) fluorescence decreases faster than the indicator’s unbinding kinetics (≥50% signal drop within ≤250 ms), or c) a 3 s.d. drop below fluorescence baseline after transient onset for ≥3 s were considered artifactual (e.g., caused by tissue motion) and excluded from further analysis. For multi-peak transients, the largest peak’s value was used for FWHM calculation. If the intensity values between peaks fell below the half-maximum value for ≥250 ms, the half-maximum point closest to the largest peak was used for FWHM calculation. If the trace’s amplitude fell below 2 s.d. above baseline for ≥400 ms, the corresponding transients were considered separate calcium spikes.

To distinguish spontaneous from evoked calcium activity, we applied additional criteria. A calcium transient was considered evoked if its onset occurred within 1 s or 5 s after stimulus- or run-onset for neurons or astrocytes with cytosolic GCaMP expression, respectively. For nuclear GCaMP expression, we allowed a 5 s time window. For all activity traces that passed the filters mentioned above, we calculated ΔF(t)/F_0_(t). F_0_ was determined with Matlab’s mode function using a 10-bin width. Traces that showed abrupt intensity changes (≥0.5 ΔF/F within ≤65 ms) were removed from the analysis. To compare neuronal and astrocyte activity, we plotted corresponding ΔF/F traces aligned on a given cell type with the other cell type being sorted based on its transient onset time.

#### Confocal imaging data

All data were processed, analyzed, and plotted using Fiji, IMARIS, and Prism software. To quantify immuno-stained tissue, we first generated digital representations of the labeled cells/structures using Imaris’ (version 9.2; Bitplane) creation wizard. Cells/structures were classified as “Spots” (to quantify the number of NeuN- and Iba-1-positive cell bodies; **Suppl. Figs. 3, 5**) or “Surfaces” (to quantify GFAP volume including cell bodies and major processes; **Suppl. Fig. 4**). For surface quantifications, we used detail and threshold levels of 0.1 and 0.4, respectively, for all tissue slices and hemispheres. These quantifications were performed for various regions of interest (ROIs) (e.g., 50 x 50 µm or 50 x 700 µm) (**Suppl. Figs. 3-5**). Volume fractions are represented in percent (**Suppl. Fig. 4b-e**).

### Analog and video data processing and analysis

All analog data were synchronously recorded at 1 kHz using DAQExpress 2.0 software (National Instruments). Analog data included the pressure sensor output from the rodent pincher system, treadmill speed, and on-off TTL signal of the miniaturized microscope‟s light source, depending on the experiment. Pinch application and mouse behavior were also recorded on a video camera (≥20 Hz; Stingray F-033, Allied Vision Technologies). To synchronize imaging with video data, we placed a near-infrared LED within the video camera‟s FOV, triggered from the microscope‟s light source drive signal (TTL pulse). Imaging and analog data were synchronized by recording the on-off TTL signal of the miniaturized microscope‟s light source together with all other analog data. To relate the animal‟s locomotor activity more closely to the imaging data, we focally restrained animals on a spherical treadmill equipped with an optical encoder (E7PD-720-118, US Digital), allowing precise readout of running speed. All analog data was processed using custom MATLAB (Mathworks) routines. The pincher and encoder traces were first cropped to the light source-on period. Pressure traces were then quantified with respect to stimulus amplitude and duration.

Encoder traces were smoothed using a sliding average (window size: 0.4 s). Locomotion onset or offset was defined as the point at which the smoothed running speed exceeded or fell below 10 mm/s. Encoder traces were analyzed concerning running speed, duration, and frequency. If the running speed fell below the 10 mm/s threshold for ≥750 ms, the local maxima were considered separate running bouts. The video data were cropped to the LED-on period. Videos were scored manually regarding pinch onset and offset. Calcium transient latency was calculated based on these measurements because video recordings provided higher temporal resolution than the pincher traces. For population analysis, data was computationally sorted using custom MATLAB (Mathworks) routines. Only trials with 1.5 ± 0.5 s pinch duration, or 3 ± 1 s run duration were included in the analysis.

### Statistical analysis

All data were analyzed and plotted using MATLAB, Excel, or GraphPad Prism software. Paired t-tests were performed to determine FOV regions with significant calcium activity (**Fig. 4f, k** and **Fig. 5c, h**), on data evaluating neuronal loss (**Suppl. Fig. 3d-f**), astrocyte activation (**Suppl. Fig. 4d-e**), microglia activation (**Suppl. Fig. 5d-e**), mouse motor performance (**Suppl. Fig. 6d-f**), hind paw sensitivity changes on the implanted side (comparing the 2-week and 4-week time points to baseline), and the potential effect of spinal plate implantation on sensory performance (**Suppl. Fig. 6b**). The latter data were collected for each hind paw before (baseline) and seven days after surgery. Two-way ANOVA was performed on data comparing hind paw sensitivity on the microprism-implanted versus the control hemisphere. All data are represented as mean ± s.e.m. unless indicated otherwise. Group sample sizes were chosen based on power analysis or previous studies. The following convention was used to indicate *P* values: ‘NS’ indicates *P*>0.05, ‘*’ indicates 0.01<*P*≤0.05, ‘**’ indicates 0.001<*P*≤0.01, ‘***’ indicates 0.0001<*P*≤0.001, and ‘****’ indicates *P*≤0.0001.

### Reporting summary

Further information on research design is available in the Research Reporting Summary linked to this paper.

### Data availability

The data that support the findings of this study will be deposited in the Brain Image Library (BIL; https://www.brainimagelibrary.org/index.html). They will also be available from the corresponding author upon reasonable request.

### Code availability

The custom Java- and Matlab-based code used to process and analyze the data will be deposited in GitHub. It will also be available from the corresponding author upon reasonable request.

## Acknowledgments

We thank members of the Nimmerjahn lab for comments on the manuscript, the Salk machine shop for technical support, and J. Chambers for mouse colony management. This work was primarily supported by the National Institutes of Health (NIH) grant R01NS108034 (A.N). It was partially supported by the NIH grants U01NS103522, U19NS112959, and U19NS123719 (A.N.), and equipment funds from C. and L. Greenfield. P.S. was supported by a Rose Hills Foundation graduate fellowship and N.A.N. by funds from an NIH T32/CMG Training Grant, Burt and Ethel Aginsky Research Scholar Award, Kavli-Helinski Endowment Graduate Fellowship, and NIH individual predoctoral fellowship (F31NS120619). The content is solely the authors’ responsibility and does not necessarily represent the official views of the NIH.

## Author contributions

P.S., E.M.C., and A.N. conceived and designed the study. P.S. developed and characterized the wearable microscopes and wrote ImageJ-based data analysis code. E.M.C. conducted the *in vivo* microprism, motor behavior, and immunostaining experiments. D.D. performed the *in vivo* multiplex imaging experiments. A. Ngo and G.G. developed Matlab- and ImageJ-based data analysis code, respectively. N.A.N. performed the sensory tests. C.L.K. produced the viral vectors. A.N. supervised the study and wrote the initial manuscript draft. All authors contributed to the text and figures, discussed the results, or provided input and edits on the manuscript.

## Supplementary Figures

**Suppl. Fig. 1.**
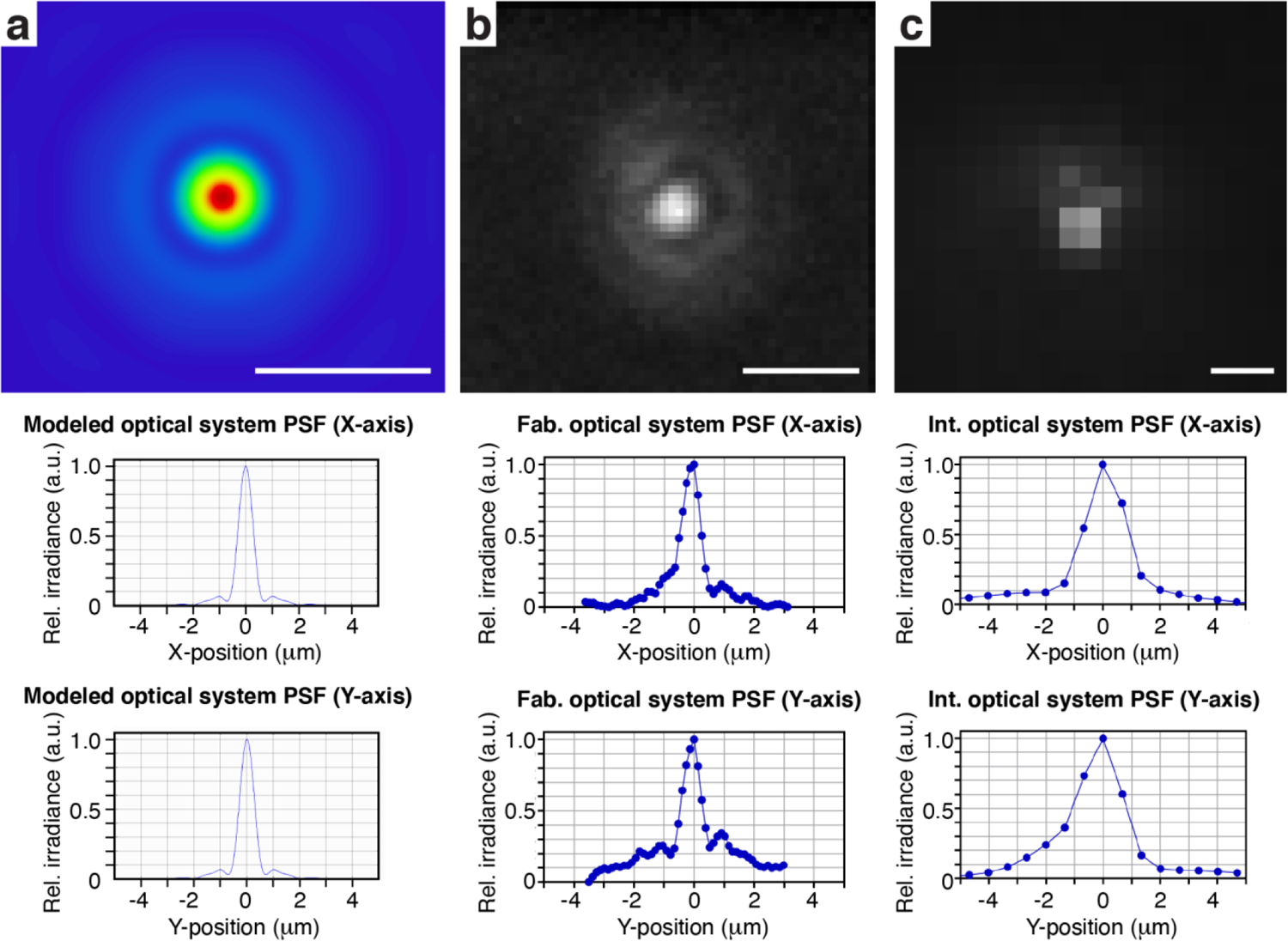
Wearable microscopes with custom-compound microlenses provide <1.5 μm lateral resolution. **a**, *Top*, the image shows the optical system’s lateral point spread function (PSF) as predicted by Zemax optical modeling. *Center*, x cross-section. *Bottom*, y cross-section. **b**, *Top*, experimentally measured lateral PSF of the fabricated optical system. *Center*, x cross-section. *Bottom*, y cross-section. **c**, *Top*, experimentally measured lateral PSF of the integrated microscope. *Center*, x cross-section. *Bottom*, y cross-section. Scale bars, 2 μm.

**Suppl. Fig. 2.**
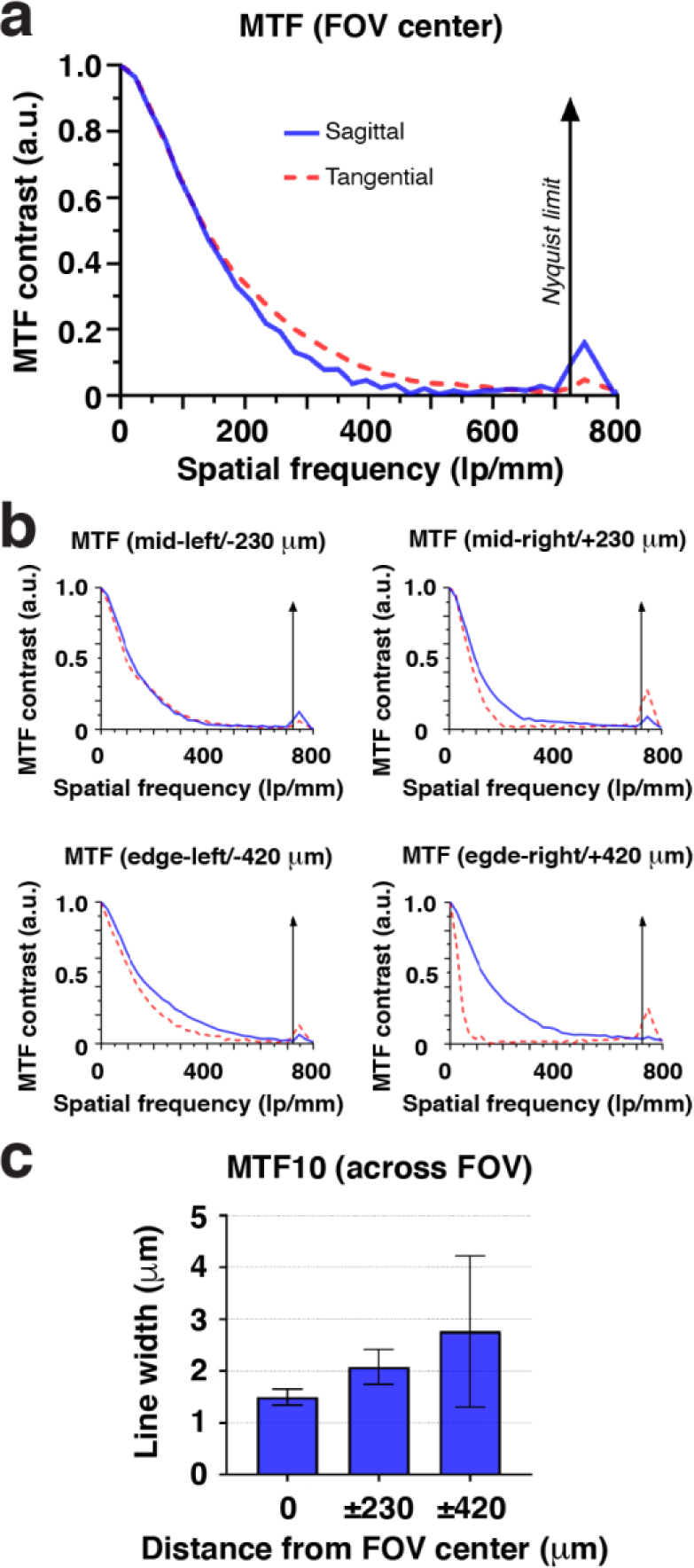
Wearable microscopes with custom-compound microlenses provide high contrast across the field of view. **a**, Modulation transfer function (MTF) of the integrated microscope measured in the center of the field of view (FOV) using the Slanted Edge test. **b**, MTF at different indicated FOV positions relative to the center. **c**, MTF contrast at 10% (MTF10) across the FOV. Displayed values are averages across similar FOV locations and horizontal and vertical Slanted Edge targets. Spatial frequencies were converted to line widths. The data are presented as mean ± s.e.m. The larger error bars toward the FOV edge likely indicate sample tilt.

**Suppl. Fig. 3.**
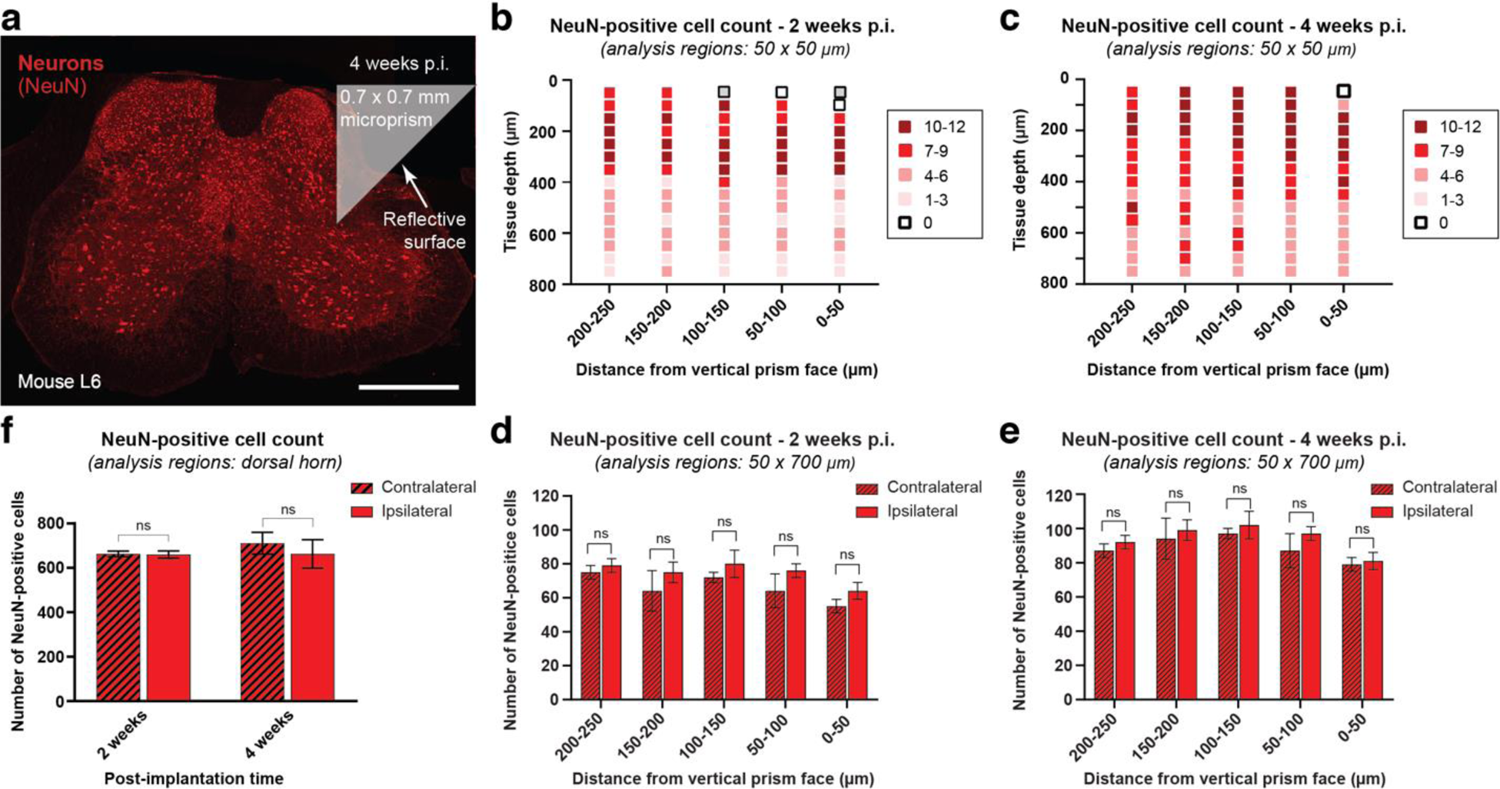
Microprism implantation does not result in a significant loss of dorsal horn neurons. **a**, Fluorescence image showing NeuN-positive cells (neurons) in a 20-μm-thick mouse spinal cord section four weeks after microprism implantation. The approximate microprism location is indicated. Scale bar, 500 μm. **b-c**, Average NeuN-positive cell count in 50 μm x 50 μm analysis regions plotted as a function of tissue depth and distance from the vertical microprism face two (b) and four weeks after microprism implantation (c). Cell density decreases with depth because of fewer cells with larger cell bodies. Depending on white matter thickness, cell density may also decrease toward the vertical microprism face. **d-e**, Average NeuN-positive cell count in 50 μm x 700 μm strips plotted as a function of distance from the vertical microprism face two (d) and four weeks after microprism implantation (e). The data are compared to equivalent regions on the non-implanted (contralateral) hemisphere. **f**, Average NeuN-positive cell count across the entire dorsal horn two and four weeks after microprism implantation. The data are compared to equivalent regions on the non-implanted (contralateral) hemisphere. The data in b-f are from 12 slices and 3 mice for each time point. Paired t-tests determined *P* values, and all data are presented as mean ± s.e.m.

**Suppl. Fig. 4.**
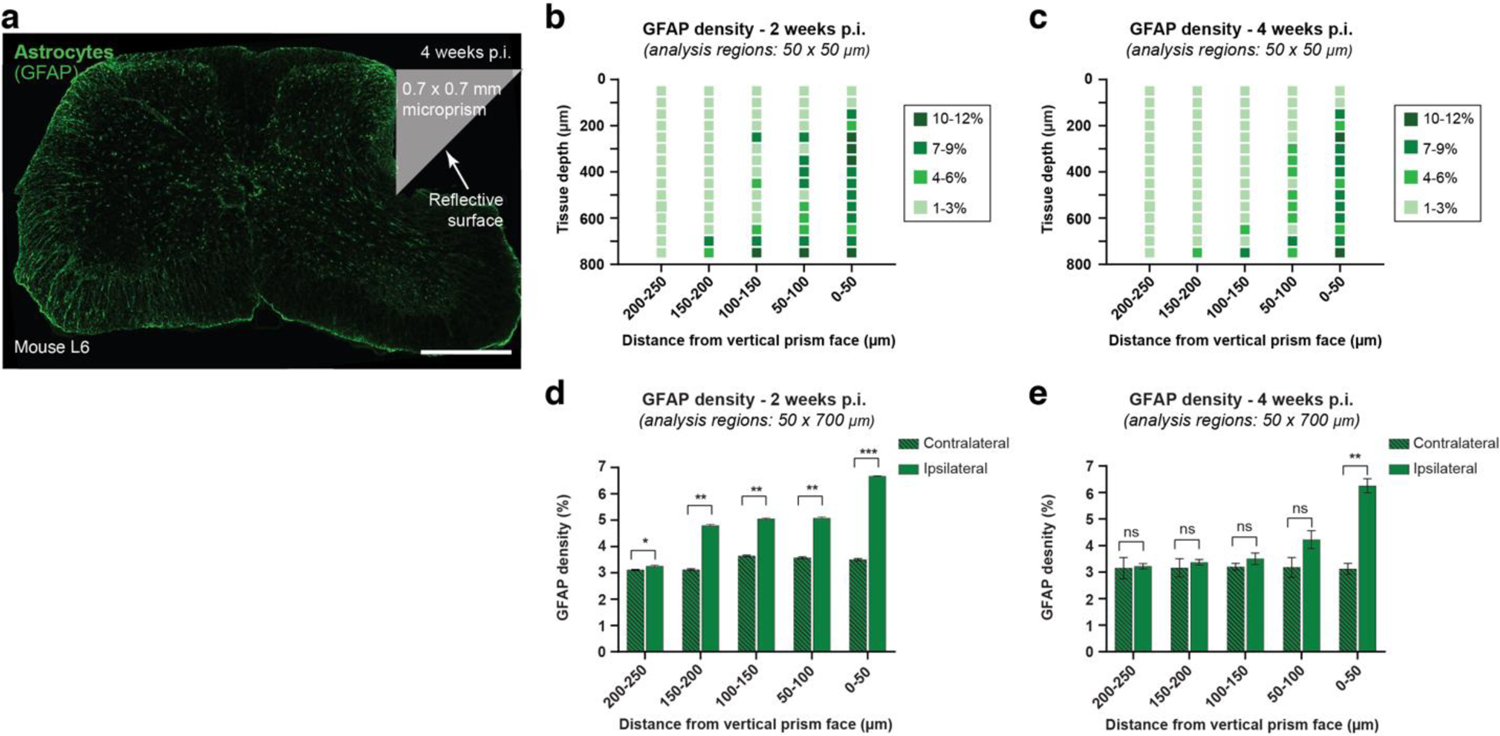
Microprism implantation leads to a transient glial fibrillary acidic protein (GFAP) upregulation near the microprism-tissue interface. **a**, Fluorescence image showing GFAP staining (astrocytes) in a 20-μm-thick mouse spinal cord section four weeks after microprism implantation. The approximate microprism location is indicated. Scale bar, 500 μm. **b-c**, Average GFAP density in 50 μm x 50 μm analysis regions plotted as a function of tissue depth and distance from the vertical microprism face two (b) and four weeks after microprism implantation (**c**). GFAP immunoreactivity decreases over time and is highly localized to regions immediately adjacent to the vertical microprism-tissue interface. d-e, Average GFAP density in 50 μm x 700 μm strips plotted as a function of distance from the vertical microprism face two (**d**) and four weeks after microprism implantation (**e**). The data are compared to equivalent regions on the non-implanted (contralateral) hemisphere. The data in b-e are from 12 slices and 3 mice for each time point. Paired t-tests determined *P* values, and all data are presented as mean ± s.e.m.

**Suppl. Fig. 5.**
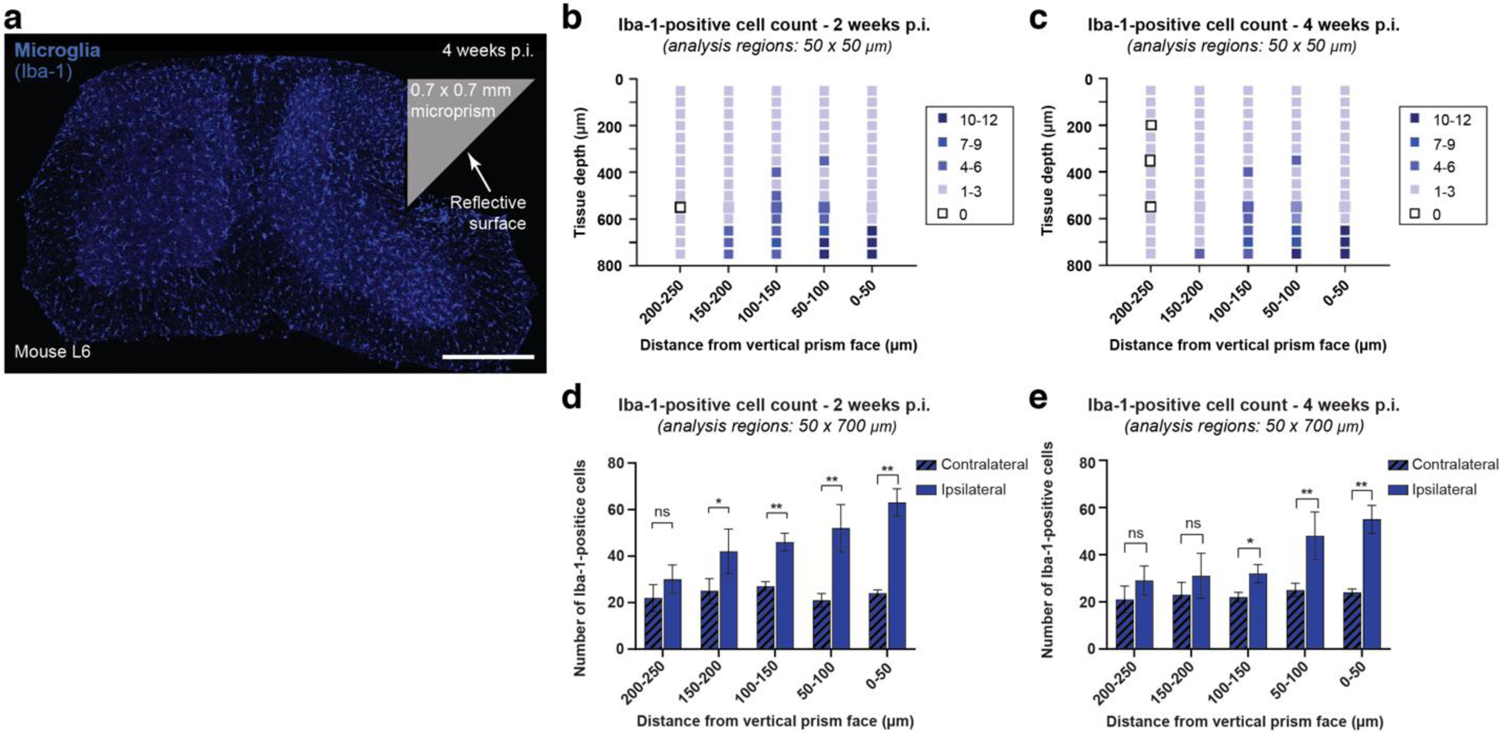
Microprism implantation leads to a transient and highly localized Iba-1 upregulation near the implanted microprism edge. **a**, Fluorescence image showing Iba-1-positive cells (microglia/macrophages) in a 20-μm-thick mouse spinal cord section four weeks after microprism implantation. The approximate microprism location is indicated. Scale bar, 500 μm. **b-c**, Average Iba-1-positive cell count in 50 μm x 50 μm analysis regions plotted as a function of tissue depth and distance from the vertical microprism face two (b) and four weeks after microprism implantation (c). Iba-1-positive cell count decreases over time and is highly localized to regions surrounding the implanted microprism edge. **d-e**, Average Iba-1-positive cell count in 50 μm x 700 μm strips plotted as a function of distance from the vertical microprism face two (d) and four weeks after microprism implantation (e). The data are compared to equivalent regions on the non-implanted (contralateral) hemisphere. The data in b-e are from 12 slices and 3 mice for each time point. Paired t-tests determined *P* values, and all data are presented as mean ± s.e.m.

**Suppl. Fig. 6.**
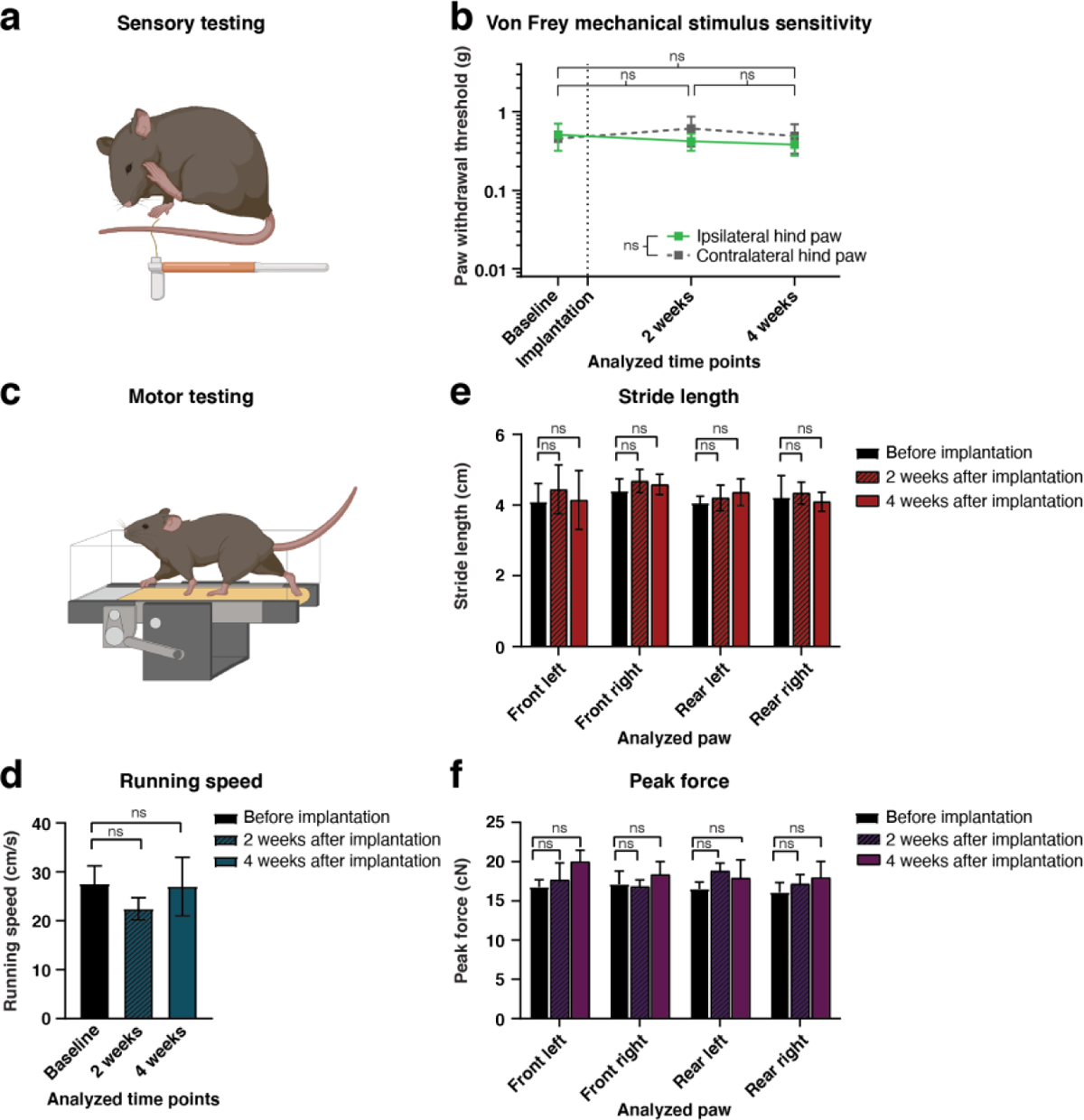
Microprism implantation does not result in overt sensory or motor deficits. **a**, Schematic showing the von Frey assay used for sensory testing. Hind paw withdrawal threshold was quantified for the implanted (ipsilateral) and non-implanted (contralateral) side before and at different time points after microprism implantation at the L4-L5 spinal level. **b**, Hind paw withdrawal threshold on the implanted (ipsilateral) and non-implanted (contralateral) side before, two, and four weeks after microprism implantation. Paw withdrawal threshold was not significantly different between time points and hemispheres (N=4 mice; ipsilateral baseline: 0.511 ± 0.193; two-week time point: 0.422 ± 0.103; four-week time point: 0.383 ± 0.108). *P* values were determined by Bonfierri-Dunn corrected paired t-tests between time points for the ipsilateral paw (*P* > 0.05) and two-way ANOVA across all time points between the contralateral and ipsilateral hind paws (*P* = 0.62). The average value of two test sessions is reported for each animal at each indicated time point (**Methods**). **c**, Schematic of the kinematic weight-bearing test used for motor testing. **d-f**, Population data showing running speed (d), front and hind paw stride length (e), and front and hind paw peak force (f) before, two, and four weeks after microprism implantation at the L4-L5 spinal level. Locomotor performance was not significantly different between time points (N=4 mice). All animals were tested at each indicated time point (**Methods**). Paired t-tests determined *P* values, and all data are presented as mean ± s.e.m.

## Supplementary Tables

**Suppl. Table 1.** Wearable microscope optical design.

| Surface | Type | Radius | Thickness | Glass | Diameter |
| --- | --- | --- | --- | --- | --- |
| OBJ | STANDARD | Infinity | 0.000000 |  | 6.000000 |
| 1 | STANDARD | Infinity | 4.500000 |  | 4.800000 |
| 2 | STANDARD | -13.65635 | 1.000000 | N-SF57 | 2.679000 |
| 3 | STANDARD | 1.758689 | 0.9479822 |  | 2.349000 |
| 4 | STANDARD | -10.38426 | 1.000000 | N-BALF5 | 2.657000 |
| 5 | STANDARD | 9.049044 | 1.1821360 | N-LASF40 | 3.988000 |
| 6 | STANDARD | -3.253426 | 0.6096153 |  | 3.785000 |
| 7 | STANDARD | Infinity | 5.000000 | FSILICA | 4.000000 |
| 8 | STANDARD | Infinity | 0.000000 |  | 4.000000 |
| 9 | STANDARD | Infinity | 0.4398171 |  | 2.000000 |
| STO | STANDARD | 5.543071 | 1.000000 | N-SF66 | 2.000000 |
| 11 | STANDARD | 1.973004 | 1.1777090 | N-PK51 | 1.9911980 |
| 12 | STANDARD | -5.628253 | 0.100000 |  | 2.994000 |
| 13 | STANDARD | 2.635428 | 1.1278330 | N-LAK34 | 3.297000 |
| 14 | STANDARD | -13.04288 | 2.000000 |  | 3.360000 |
| 15 | STANDARD | Infinity | 0.100000 | D263M | 0.8690751 |
| IMA | STANDARD | Infinity |  |  | 0.8293949 |

## Supplementary Movies

**Suppl. Movie 1 High-speed translaminar imaging of tail pinch-evoked calcium activity in behaving GFAP-GCaMP6f mice**. Example time-lapse recording acquired with the wearable microscope showing noxious tail pinch-evoked calcium excitation in spinal astrocytes. The data were obtained at ∼42.3 fps and ∼75 μm focal depth from the vertical tissue-microprism interface four weeks after microprism implantation. Elapsed time is indicated in the upper right corner (total duration: 30 s). To allow a precise readout of locomotor activity, the mouse was placed on a spherical treadmill. Noxious pinch triggered coordinated calcium excitation in the central field of view regions corresponding to upper dorsal horn laminae. Scale bar, 100 μm.

**Suppl. Movie 2 High-speed translaminar imaging of motor-evoked calcium activity in behaving GFAP-GCaMP6f mice**. Example time-lapse recording acquired with the wearable microscope showing running-evoked calcium excitation in spinal astrocytes. The data were obtained at ∼40.3 fps and ∼75 μm focal depth from the vertical tissue-microprism interface four weeks after microprism implantation. Elapsed time is indicated in the upper right corner (total duration: 1 min 19 s). To allow a precise readout of locomotor activity, the mouse was placed on a spherical treadmill. Running triggered coordinated calcium excitation in the lower field of view regions likely corresponding to spinal premotor areas. Scale bar, 100 μm.

**Suppl. Movie 3 High-speed translaminar imaging of tail pinch-evoked calcium activity in behaving Tac1-GCaMP6f mice**. Example time-lapse recording acquired with the wearable microscope showing noxious tail pinch-evoked calcium excitation in spinal Tac1-expressing neurons. The data were obtained at ∼43.2 fps and ∼75 μm focal depth from the vertical tissue-microprism interface four weeks after microprism implantation. Elapsed time is indicated in the upper right corner (total duration: 33 s). To allow a precise readout of locomotor activity, the mouse was placed on a spherical treadmill. Noxious pinch triggered coordinated calcium excitation in the central field of view regions corresponding to upper dorsal horn laminae. Scale bar, 100 μm.

## References

1. Koch, S. C., Acton, D. & Goulding, M. Spinal circuits for touch, pain, and itch. Annu Rev Physiol 80, 189–217, doi:10.1146/annurev-physiol-022516-034303 (2018).

2. Abraira, V. E. & Ginty, D. D. The sensory neurons of touch. Neuron 79, 618–639, doi:10.1016/j.neuron.2013.07.051 (2013).

3. Peirs, C. & Seal, R. P. Neural circuits for pain: Recent advances and current views. Science 354, 578–584, doi:10.1126/science.aaf8933 (2016).

4. Sekiguchi, K. J. et al. Imaging large-scale cellular activity in spinal cord of freely behaving mice. Nat Commun 7, 11450, doi:10.1038/ncomms11450 (2016).

5. Kronschlager, M. T. et al. Gliogenic LTP spreads widely in nociceptive pathways. Science 354, 1144–1148, doi:10.1126/science.aah5715 (2016).

6. Kelley, K. W. et al. Kir4.1-dependent astrocyte-fast motor neuron interactions are required for peak strength. Neuron 98, 306–319 e307, doi:10.1016/j.neuron.2018.03.010 (2018).

7. Kohro, Y. et al. Spinal astrocytes in superficial laminae gate brainstem descending control of mechanosensory hypersensitivity. Nat Neurosci 23, 1376–1387, doi:10.1038/s41593-020-00713-4 (2020).

8. Xu, Q. et al. Astrocytes contribute to pain gating in the spinal cord. Sci Adv 7, eabi6287, doi:10.1126/sciadv.abi6287 (2021).

9. Nelson, N. A., Wang, X., Cook, D., Carey, E. M. & Nimmerjahn, A. Imaging spinal cord activity in behaving animals. Exp Neurol 320, 112974, doi:10.1016/j.expneurol.2019.112974 (2019).

10. Cheng, Y. T., Lett, K. M. & Schaffer, C. B. Surgical preparations, labeling strategies, and optical techniques for cell-resolved, in vivo imaging in the mouse spinal cord. Exp Neurol 318, 192–204, doi:10.1016/j.expneurol.2019.05.010 (2019).

11. Andermann, M. L. et al. Chronic cellular imaging of entire cortical columns in awake mice using microprisms. Neuron 80, 900–913, doi:10.1016/j.neuron.2013.07.052 (2013).

12. Low, R. J., Gu, Y. & Tank, D. W. Cellular resolution optical access to brain regions in fissures: imaging medial prefrontal cortex and grid cells in entorhinal cortex. Proc Natl Acad Sci U S A 111, 18739–18744, doi:10.1073/pnas.1421753111 (2014).

13. Heys, J. G., Rangarajan, K. V. & Dombeck, D. A. The functional micro-organization of grid cells revealed by cellular-resolution imaging. Neuron 84, 1079–1090, doi:10.1016/j.neuron.2014.10.048 (2014).

14. Aharoni, D. & Hoogland, T. M. Circuit investigations with open-source miniaturized microscopes: Past, present and future. Front Cell Neurosci 13, 141, doi:10.3389/fncel.2019.00141 (2019).

15. Chen, S. et al. Miniature fluorescence microscopy for imaging brain activity in freely-behaving animals. Neurosci Bull 36, 1182–1190, doi:10.1007/s12264-020-00561-z (2020).

16. Matz, G., Messerschmidt, B. & Gross, H. Design and evaluation of new color-corrected rigid endomicroscopic high NA GRIN-objectives with a sub-micron resolution and large field of view. Opt Express 24, 10987–11001, doi:10.1364/OE.24.010987 (2016).

17. Simonetti, M. et al. Nuclear calcium signaling in spinal neurons drives a genomic program required for persistent inflammatory pain. Neuron 77, 43–57, doi:10.1016/j.neuron.2012.10.037 (2013).

18. Nimmerjahn, A., Mukamel, E. A. & Schnitzer, M. J. Motor behavior activates Bergmann glial networks. Neuron 62, 400–412, doi:10.1016/j.neuron.2009.03.019 (2009).

19. Merten, K., Folk, R. W., Duarte, D. & Nimmerjahn, A. Astrocytes encode complex behaviorally relevant information. bioRxiv, doi: https://doi.org/10.1101/2021.1110.1109.463784 (2021).

20. Polgar, E. et al. Substance P-expressing neurons in the superficial dorsal horn of the mouse spinal cord: Insights into their functions and their roles in synaptic circuits. Neuroscience 450, 113–125, doi:10.1016/j.neuroscience.2020.06.038 (2020).

21. Huang, T. et al. Identifying the pathways required for coping behaviours associated with sustained pain. Nature 565, 86–90, doi:10.1038/s41586-018-0793-8 (2019).

22. Flusberg, B. A. et al. High-speed, miniaturized fluorescence microscopy in freely moving mice. Nat Methods 5, 935–938, doi:10.1038/nmeth.1256 (2008).

23. Ghosh, K. K. et al. Miniaturized integration of a fluorescence microscope. Nat Methods 8, 871–878, doi:10.1038/nmeth.1694 (2011).

24. Kronschlager, M. T. et al. Lamina-specific properties of spinal astrocytes. Glia 69, 1749–1766, doi:10.1002/glia.23990 (2021).

25. Molofsky, A. V. et al. Astrocyte-encoded positional cues maintain sensorimotor circuit integrity. Nature 509, 189–194, doi:10.1038/nature13161 (2014).

26. Harrison, M. et al. Vertebral landmarks for the identification of spinal cord segments in the mouse. Neuroimage 68, 22–29, doi:10.1016/j.neuroimage.2012.11.048 (2013).

27. Ceto, S., Sekiguchi, K. J., Takashima, Y., Nimmerjahn, A. & Tuszynski, M. H. Neural stem cell grafts form extensive synaptic networks that integrate with host circuits after spinal cord injury. Cell Stem Cell 27, 430–440 e435, doi:10.1016/j.stem.2020.07.007 (2020).

28. Sabatini, B. L. & Tian, L. Imaging neurotransmitter and neuromodulator dynamics in vivo with genetically encoded indicators. Neuron 108, 17–32, doi:10.1016/j.neuron.2020.09.036 (2020).

## References

29. Garcia, A. D., Doan, N. B., Imura, T., Bush, T. G. & Sofroniew, M. V. GFAP-expressing progenitors are the principal source of constitutive neurogenesis in adult mouse forebrain. Nat Neurosci 7, 1233–1241, doi:10.1038/nn1340 (2004).

30. Madisen, L. et al. A toolbox of Cre-dependent optogenetic transgenic mice for light-induced activation and silencing. Nat Neurosci 15, 793–802, doi:10.1038/nn.3078 (2012).

31. Harris, J. A. et al. Anatomical characterization of Cre driver mice for neural circuit mapping and manipulation. Front Neural Circuits 8, 76, doi:10.3389/fncir.2014.00076 (2014).

32. Bonin, R. P., Bories, C. & De Koninck, Y. A simplified up-down method (SUDO) for measuring mechanical nociception in rodents using von Frey filaments. Mol Pain 10, 26, doi:10.1186/1744-8069-10-26 (2014).

33. Dubbs, A., Guevara, J. & Yuste, R. moco: Fast motion correction for calcium imaging. Front Neuroinform 10, 6, doi:10.3389/fninf.2016.00006 (2016).

34. Patriarchi, T. et al. Ultrafast neuronal imaging of dopamine dynamics with designed genetically encoded sensors. Science 360, doi:10.1126/science.aat4422 (2018).

